# Interpretable abstractions of artificial neural networks predict behavior and neural activity during human information gathering

**DOI:** 10.1101/2025.06.24.661282

**Authors:** Simone D’Ambrogio, Jan Grohn, Nima Khalighinejad, Marcelo Mattar, Laurence Hunt, Matthew F.S. Rushworth

## Abstract

It has been suggested that humans and other animals are driven by a fundamental desire to acquire information about opportunities available in their environments. Not only might such a desire explain pathological behaviors, but it may be needed to account for how everyday decisions are resolved. Here, we combine artificial neural networks (ANNs) with symbolic regression to extract an expressive yet interpretable model that specifies how human participants evaluate decision-relevant information during choice. This model accounts for behavior in our own data and in previous work, outperforming existing accounts of information sampling such as the Upper Confidence Bound heuristic. This modelling approach has broad potential for uncovering novel patterns in behavior and cognitive processes, while also specifying them in human-interpretable formats. We then used the value of information derived by our model, together with ultra-high field neuroimaging, to examine activity across a suite of subcortical neuromodulatory nuclei and two cortical regions that influence these nuclei. This established roles for midbrain dopaminergic nuclei, anterior cingulate cortex, and anterior insula in mediating the influence of value of information on behavior.

## Introduction

Decisions should be based on evidence. Once sufficient evidence has been sampled, the agent can decide which option to select (Fig. 1a, top). But in addition to guiding the choice, evidence should also simultaneously be used to evaluate whether there is sufficient information to warrant a decision, or whether further information must be sampled (Fig. 1a, bottom) ^1–4^. When sampling information, attentional constraints mean that decision makers typically only focus on one option at a time ^5^. There is therefore a further decision, as to whether more evidence should be gathered from the currently attended option or whether attention should be shifted to an alternative (Fig. 1a, bottom). Gathering more information can improve decision quality, however it also comes at the expense of time, energy, and lost opportunities to engage in other activities. Therefore, knowing when to stop learning from the environment and use the acquired knowledge to make a choice is crucial for effective decision-making. An illustration of this dilemma is found in the 14th-century thought experiment associated with Jean Buridan ^6^ but discussed by many others before and since. In the scenario an ass stands exactly midway between two identical piles of hay (Fig. 1a). In this extreme example, the piles are so similar to one another that the ass cannot choose between them. The ass contemplates the piles searching for evidence that one is better than the other but, without a mechanism to stop deliberating, the ass fails to make a choice and is left without anything. While several studies have shed light on the computational and neural mechanisms underlying value learning and option selection ^7,8^, the factors that determine when to stop gathering information and commit to a final decision, as well as which options to sample from, remain an active area of investigation ^1,4,9,10^.

**Fig. 1.**
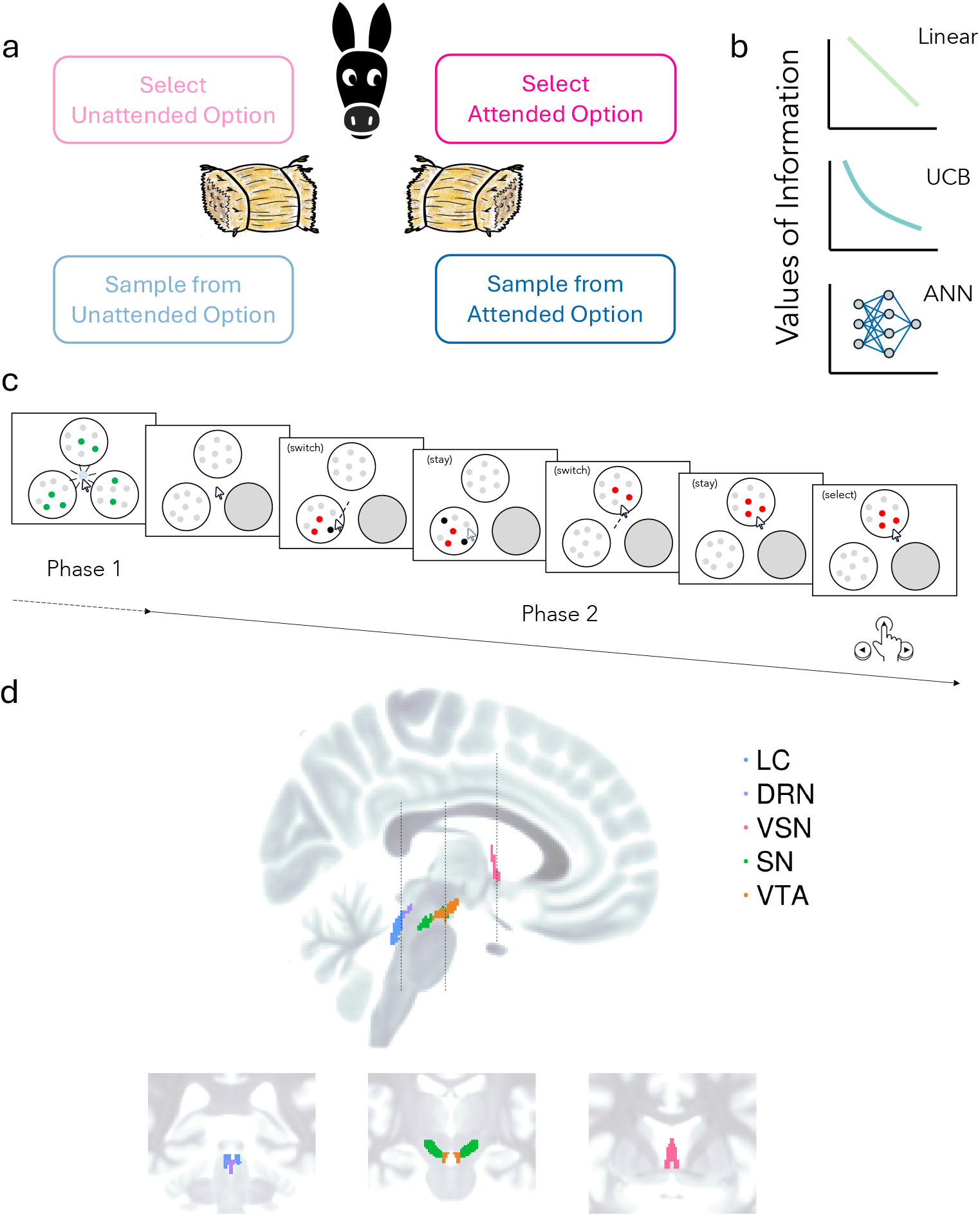
Information sampling task. a, The decision-making process involves two key decisions: whether to gather more information or make a selection (top vs. bottom), and if gathering information, whether to sample from the currently attended option or switch to the alternative (top right vs. top left). b, Three approaches to computing the value of information: a linear function that assigns constant value to additional information (top), Upper Confidence Bound (UCB) algorithm that captures diminishing returns (middle), and an artificial neural network (ANN) that learns the optimal mapping between state variables and information value (bottom). c, Task structure showing the two phases of the information sampling task. In Phase 1, participants are presented with three patches of dots covered by green or grey covers. After revealing the green-covered dots in each patch, one patch is blocked (grey circle). In Phase 2, participants can freely sample information by hovering over patches, with grey-covered dots revealing their true colors sequentially, before making a final selection. d, Brain regions of interest: locus coeruleus (LC), dorsal raphe nucleus (DRN), ventral septal nucleus (VSN), substantia nigra (SN), and ventral tegmental area (VTA), which have been implicated in uncertainty processing and information sampling.

A common approach in cognitive neuroscience involves formulating hypotheses about the computations that drive specific behaviors and formalizing these through mathematical models. These models serve as explicit computational representations of cognitive processes, allowing researchers to generate quantitative predictions about behavioral and neural data. In line with this framework, two hypotheses can be considered to explain how individuals guide their information sampling behavior. The first posits that people compute the value of gathering additional information as a linear function of an option’s uncertainty (e.g., the amount of missing knowledge; Fig. 1b top) ^11–13^. This approach would prioritize further sampling from more uncertain options. However, theories of sequential sampling processes and Bayesian updating indicate that repeatedly sampling from the same option yields diminishing returns (Fig. 1b, middle), motivating a second hypothesis: that the value of information is computed using a non-linear function of uncertainty ^1,9^. The Upper Confidence Bound (UCB) algorithm, a widely used exploration heuristic that captures this non-linear relation, can formalize this hypothesis. Both linear and UCB models aim to characterise the functional form of a cognitive computation, such as the value of sampling, in a psychologically interpretable way.

While the concept of diminishing returns helps narrow down the space of hypotheses, there is still a wide range of non-linear functions that could describe how people value information. This presents a challenge in psychology and neuroscience, where it is often difficult to select a specific instantiation of a general principle, slowing the pace of new discoveries. An alternative approach is to use machine learning to provide a more flexible, data-driven model of participants’ choices – for example, using artificial neural networks (ANNs) to fit behaviour. ANNs can learn complex mappings from data, vastly expanding the space of candidate functions that can be considered (Fig. 1b, bottom) ^14,15^. The universal function approximation theorem underpins this method, asserting that sufficiently deep neural networks can model any continuous function to arbitrary precision, given sufficient training data. This capability allows us to learn directly from data how individuals might compute the value of information to guide their behavior, without needing to specify the exact form of the function in advance. By comparing an ANN’s performance against established, fixed-form functions (like linear or UCB), we can directly assess whether these more constrained models adequately capture the complexities of information valuation. If an ANN provides a better account of behavior, it suggests that the underlying computations are more nuanced than these assumed by the simpler benchmarks. ANNs can thus be uses as a tool to discover a potentially more accurate functional description of how people assign value to information. However, this expressive power comes at a cost. While ANNs may yield more accurate predictions, they typically lack interpretability. The learned representations are distributed, high-dimensional, and opaque, making it difficult to extract mechanistic or symbolic insights about underlying cognitive processes.

Here, we propose a modelling approach that yields expressive yet interpretable models of choice. First, we integrate data-driven and knowledge-driven components into a hybrid model of subjects’ information sampling and decision making behavior ^14^. The knowledge-driven component incorporates established cognitive principles, such as the mechanisms underlying option selection ^8,16^. The data-driven component utilizes ANNs to model aspects of cognition that are difficult to define, such as complex information sampling strategies. The hybrid model can be more expressive than a fully knowledge-driven approach, while also being more interpretable and data-efficient than a fully data-driven approach. Second, we additionally apply symbolic regression ^17,18^ to the trained ANN part of the model to recover a compact, four-parameter approximation to the ANN’s learned function. This final step enables data-driven discovery of novel, fully interpretable process-level or symbolic models of cognition, offering a new route for theory generation in psychology and neuroscience. We show that this newly recovered model generalises to an entirely separate experimental dataset on human information sampling ^19^.

We used our approach to model human behavioral data from a task designed to assess evaluation of information and choice selection (Fig. 1c). Several brain regions are thought to play key roles in information sampling under uncertainty. Notably all the neuromodulatory systems, with their origins in the ventral tegmental area (VTA), substantia nigra (SN), dorsal raphe nucleus (DRN), locus coeruleus (LC), and ventral septal nucleus (VSN) have at one time or another been proposed to reflect uncertainty or the potential for information gain ^3,20–26^(Fig. 1d). It is less clear whether each neuromodulatory system has a specific or unique relationship with uncertainty. It has been difficult to record simultaneously from multiple nuclei in animal models, especially in the same individual. At the same time, it has been difficult to identify reliable signals from some of these nuclei in humans using standard neuroimaging approaches which have limited spatial resolution. Here, we exploit high resolution (1mm isotropic), rapid repetition time (1.378 s), accelerated ultra-high field (7T) functional magnetic resonance imaging together with an established suite of careful data collection and preprocessing steps to measure activity in VTA, SN, DRN, LC, and VSN simultaneously ^26,27^. We positioned our functional magnetic resonance imaging (fMRI) data collection to record simultaneously from all five ascending neuromodulatory nuclei and two interconnected cortical areas – the anterior insula (AI), and anterior cingulate cortex (ACC) - which are distinguished by projecting to these nuclei and exhibiting related activity in other contexts ^3,21,28–32^. Guided by the hybrid ANN model, we identified specific and distinct patterns of activity across these seven brain areas, linked to distinct components of information valuation and evidence accumulation for action selection.

## Results

### Sampling Behavior Adaptively Scales with Task Difficulty and Uncertainty

Twenty participants completed an information-sampling task (Fig. 1c) inside a 7T MRI scanner. In each trial, they were presented with two patches of 100 moving dots. Participants were informed that the true color of each dot was either red or black, and the goal was to select the patch with the highest number of red dots. At the start of each trial, however, the true color of the dots in each patch was unknown because they were hidden under green or grey covers. The number of green dot covers (revealed simultaneously upon first hover) varied from 5 to 30, and the grey dot covers (revealed sequentially) correspondingly ranged from 70 to 95. Participants used a trackball to hover over a patch and this led to the true colors (red or black) being gradually revealed.

Upon hovering, participants had to wait 2 seconds before any new information appeared. After the waiting period, the green dot covers were all removed to reveal which dots were either red or black. The grey covers of individual dots were then also removed, but they were removed one by one every 150ms, again revealing either a red or black dot. Each patch, therefore, provided a different amount of initial information on sampling, indicated by green dots, which varied between trials. For instance, if a patch had 20 green dots, the color of these dots would be revealed as red or black during the first sample of the first visit to that patch. The grey dots in the patch were then revealed as either red or black one-by-one. By manipulating the amount of initial information gain, this design allowed us to separate the time spent in a patch from the uncertainty about the number of red dots in that patch.

This design also allowed us to examine how background uncertainty, as well as the uncertainty of the options themselves, affects sampling behavior. By signaling the initial amount of information with green dots, participants could estimate the starting uncertainty of each option. Once all green dots were shown, one of the three options was blocked, leaving only two options available. The blocked option’s uncertainty (background uncertainty) was unaffected by participants’ actions and did not impact the task of selecting the patch with more red dots. Each participant completed four sessions, each of which lasted for 25 minutes. Participants were instructed and incentivized to make as many correct choices as possible in each session. Each correct choice yielded one point. Notably, because each session had a fixed duration, spending excessive time revealing all dots in a single trial reduced the number of subsequent trials participants could attempt, thereby limiting their overall potential rewards.

Participants exhibited varying preferences for speed versus accuracy in their sampling behavior (Fig. 2a). Some participants opted to spend more time gathering information, aiming to increase accuracy, while others prioritized faster decisions, accepting a higher risk of error (correlation between amount of information and accuracy: *r* = 0.74, *t* = 4.33, *P* = 4.00 × 10^−4^). As noted, each trial differed in two key aspects: the initial uncertainty of each option, as indicated by the green dots, and the final difference in red dots between the patches, ranging from a difference of 30 (easy trials) to 10 (difficult trials). Participants tended to gather more samples when the initial uncertainty was higher (there were fewer green dots, which were uncovered at the beginning of the sampling period, and more grey dots which were only uncovered one-by-one during sampling: *β* = −0.069, *s*.*e*. = 0.013, *z* = −5.18, *P* = 2.27 × 10^−7^) and when the final discrepancy in red dots was smaller (*β* = −0.131, *s*.*e*. = 0.019, *z* = −6.87, *P* = 6.48 × 10^−12^; Fig. 2b). To assess whether this pattern is adaptive we estimated the optimal policy for this task solving the Bellman equation using dynamic programming ^33,34^ (see Methods). We simulated action sequences from this optimal agent and compared them to the actual action sequences exhibited by participants. We found that participants’ sampling behavior closely matched the optimal policy derived from our computational analysis: they adaptively increased their sampling time when initial uncertainty (100 – number of green dots, or in other words initial number of grey dots) was higher or when decisions were more challenging (lower disparity between the number of red dots associated with each option). Some participants deviated from this policy by sampling more information than was optimal - but while their accuracy generally increased, they tended ultimately to earn fewer points (Fig. 2a) due to the opportunity cost of performing more trials.

**Fig. 2.**
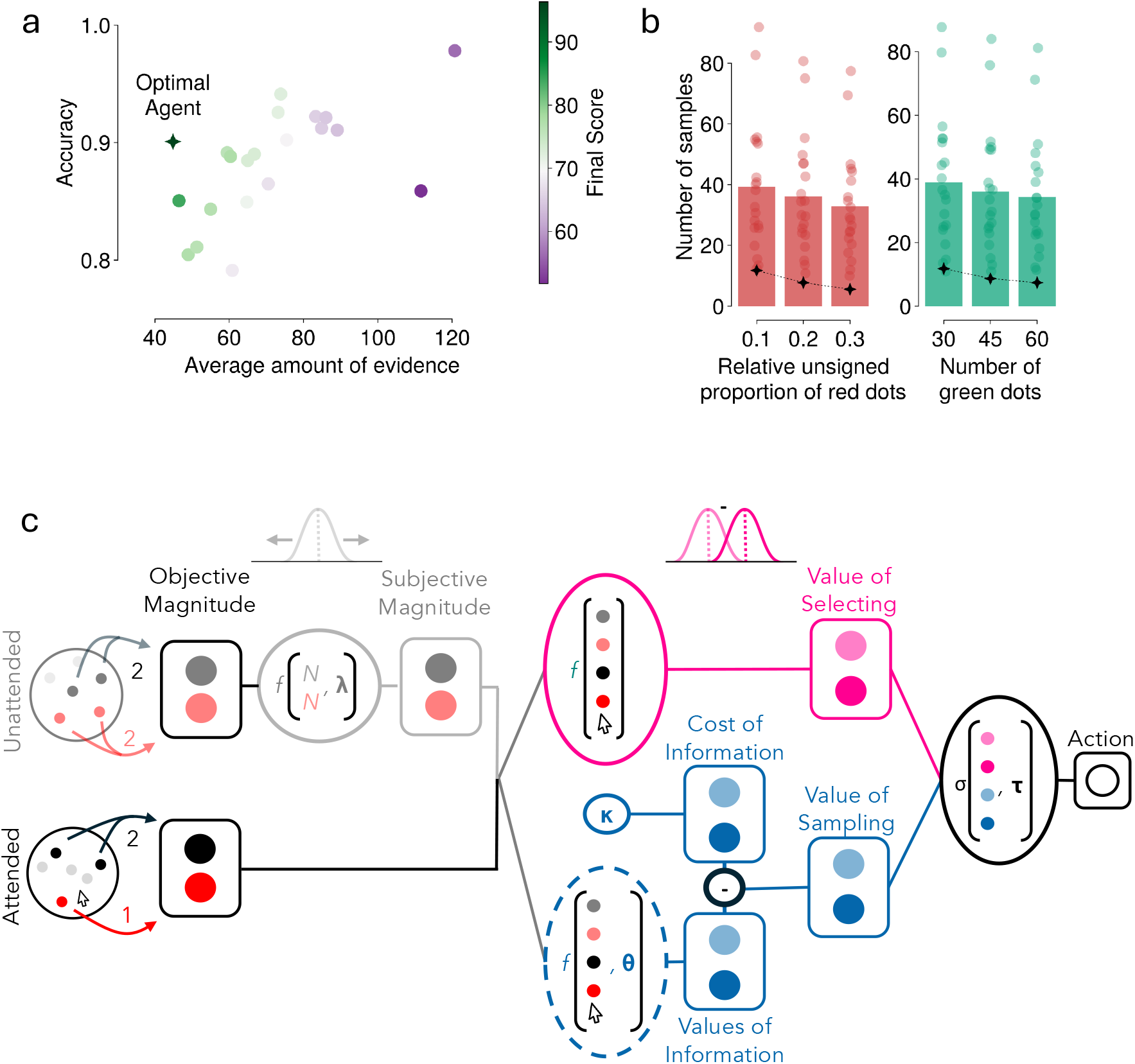
Sampling behavior and computational model. a, Relationship between accuracy and average amount of evidence collected. Each colored dot represents an individual participant, with color indicating their final score according to the color scale on the right. The dark green star marks the position of the optimal agent. b, Number of samples collected under different task conditions. Left: Bar graph showing the number of samples collected across three levels of relative unsigned proportion of red dots between patches (0.1, 0.2, 0.3). Right: Bar graph showing the number of samples collected across three levels of initial information (30, 45, 60 green dots). Individual participant data points are shown as colored dots, and black stars indicate the optimal agent’s behavior. c, Schematic of the computational model. The model transforms objective magnitudes (number and color of dots in each patch) into subjective magnitudes through attentional discounting and memory decay functions. These subjective magnitudes are used to compute the value of selecting each patch (pink) and the value of sampling more information (blue), which together determine the final action. The dotted blue circle highlights the component that computes the value of information, which is implemented using either a linear function, UCB algorithm, or artificial neural network.

Task difficulty was determined partly by the difference between patches in terms of the proportion of red dots (i.e. total number of red dots divided by 100); when this difference in proportions was smaller, the task was more difficult. However, the task was also more challenging when the mean of the two proportions were closer to 0.5, because the posterior belief about the proportion of red dots (which can be modeled using a Beta distribution) is wider (there is more uncertainty in the estimate) and so there is more overlap between the distributions associated with the two patches, when their mean is near 0.5, given the same difference in proportion of red dots. Conversely, when the mean proportion is closer to the boundaries of 0 or 1, the posterior belief is narrower, making it easier to distinguish the two estimates from one another. For example, if 10 dots are revealed in each patch, discriminating between one patch with 4 red dots and another with 6 red dots (a difference in proportion of 0.2, mean proportion 0.5) is harder than discriminating between a patch with 1 red dot and another with 3 red dots (also a difference of 0.2, but mean proportion 0.2). We tested whether participants adapted their sampling strategy based on the inherent difficulty of discriminating proportions near 0.5 by examining how sampling behavior varied with the absolute distance of the mean proportion from Both human participants and the optimal agent gathered more samples when the mean proportions were closer to 0.5 (participants: *β* = −0.085, *s*.*e*. = 0.017, *z* = −4.87, *P* = 1.09 × 10^−6^; optimal agent: *β* = −0.038, *s*.*e*. = 0.015, *z* = −2.44, *P* = 0.015), after controlling for the total number of revealed dots and the relative difference between patches. This pattern suggests that participants adapted their sampling strategy based on the inherent difficulty of the task.

Finally, we examined which kinds of uncertainty influenced participants’ sampling behavior. In our task design, participants knew the initial uncertainty (number of green dots) of the blocked option, which should be irrelevant for optimal sampling between the two available options, as well as the uncertainty of the decision-relevant options that were available to be chosen. We tested whether this irrelevant uncertainty affected participants’ sampling behavior using a mixed-effects Poisson regression model. It did not. The analysis revealed that participants sampled less from the attended option when this attended option had more initial green dots (*β* = −0.138, *s*.*e*. = −0.017, *z* = −8.11, *P* = 5.24 × 10^−16^). They also sampled more from the attended option when the other option, the unattended option, had more initial green dots (*β* = 0.173, *s*.*e*. = 0.024, *z* = 7.19, *P* = 6.42 × 10^−13^). Crucially, however, the number of green dots in the blocked option did not significantly influence sampling behavior (*β* = −0.006, *s*.*e*. = 0.006, *z* = −1.01, *P* = 0.310), suggesting that participants were able to appropriately ignore task-irrelevant background uncertainty even when their behavior was influenced by task-relevant uncertainty. This pattern held true even immediately after exposure to the blocked option (no significant interaction between blocked dots and first visit, *β* = 0.011, *s*.*e*. = 0.008, *z* = 1.41, *P* = 0.158) and remained consistent across sessions, indicating that participants maintained this optimal strategy throughout the experiment. Consequently, this background uncertainty will not be the focus of our subsequent modeling and neural analyses. These results suggest that participants were able to distinguish between relevant and irrelevant sources of uncertainty in their sampling behavior.

### The ANN-derived value of information predicts participants’ sampling decisions

To investigate participants’ strategies, we developed and fitted a computational model that takes as input the current state (i.e. the number of red and black dots in both patches) to compute action-specific values. We considered four actions: staying (sampling from the currently attended patch; Fig. 2c; Fig. 1a, bottom right), switching (sampling from the unattended patch; Fig. 2c; Fig. 1a, bottom left), and selecting either the attended or unattended patch (Fig. 1a top; Fig. 2c). To make any one of these four actions, the model maintains and updates its beliefs about the likely number of red dots in both the currently attended patch and the unattended patch. For the attended patch, this belief is updated directly based on the colors of the dots revealed during ongoing sampling.

For the unattended patch, where there is no direct sensory input, the model relies on information presumed to be stored in working memory. The representation of this information is handled differently depending on whether the patch has been previously sampled in the trial: if the unattended patch has not yet been visited in the trial, its initial state (and thus the potential value of sampling it) is estimated using the number of green dots. The number of green dots provides a measure of the information that would be gained upon a first visit, influencing the model’s assessment of its current uncertainty. To estimate the proportion of red dots within it, the model draws upon information from the currently attended patch, effectively using the ongoing sampling as a cue. If the unattended patch has been visited previously, the model relies on its memory of the dots observed during those past visits. However, this remembered information is assumed to degrade the longer the patch remains unattended, leading to an increase in uncertainty about its contents over time^35,36^. The detailed implementation of these transformations is described in the Methods.

A critical aspect that makes effective information sampling possible is evaluating the marginal value of acquiring additional information from each option. To evaluate how participants computed this value of information, we compared three distinct models with cross validation. The first model assumes a linear relationship between the value of information and the number of collected samples (the total number of red and black dots revealed after removal of the green or grey covers). This linear model assigns the same value to new information, regardless of the number of previously collected samples (Fig. 3a, top). In other words, each new sample reduces the value of learning about the proportion of red dots in the patch by the same amount. The second model implements an Upper Confidence Bound (UCB) function to estimate the value of information. This model assigns decreasing value to information as the number of collected samples increases (Fig. 3a, middle). Finally, the third model uses an artificial neural network (ANN) to learn from the data the mapping between the state variables (such as the number and color of revealed dots) and the value of gathering additional information (Fig. 3a, bottom). This approach, which we refer to as a hybrid model, combines the data-driven nature of the ANN with the theory-driven design of the cognitive model in which the ANN is instantiated. The advantage of this approach is that it gives us the flexibility to capture potentially complex, non-linear relationships between task variables and information value without making strong a priori assumptions about the functional form of this relationship.

**Fig. 3.**
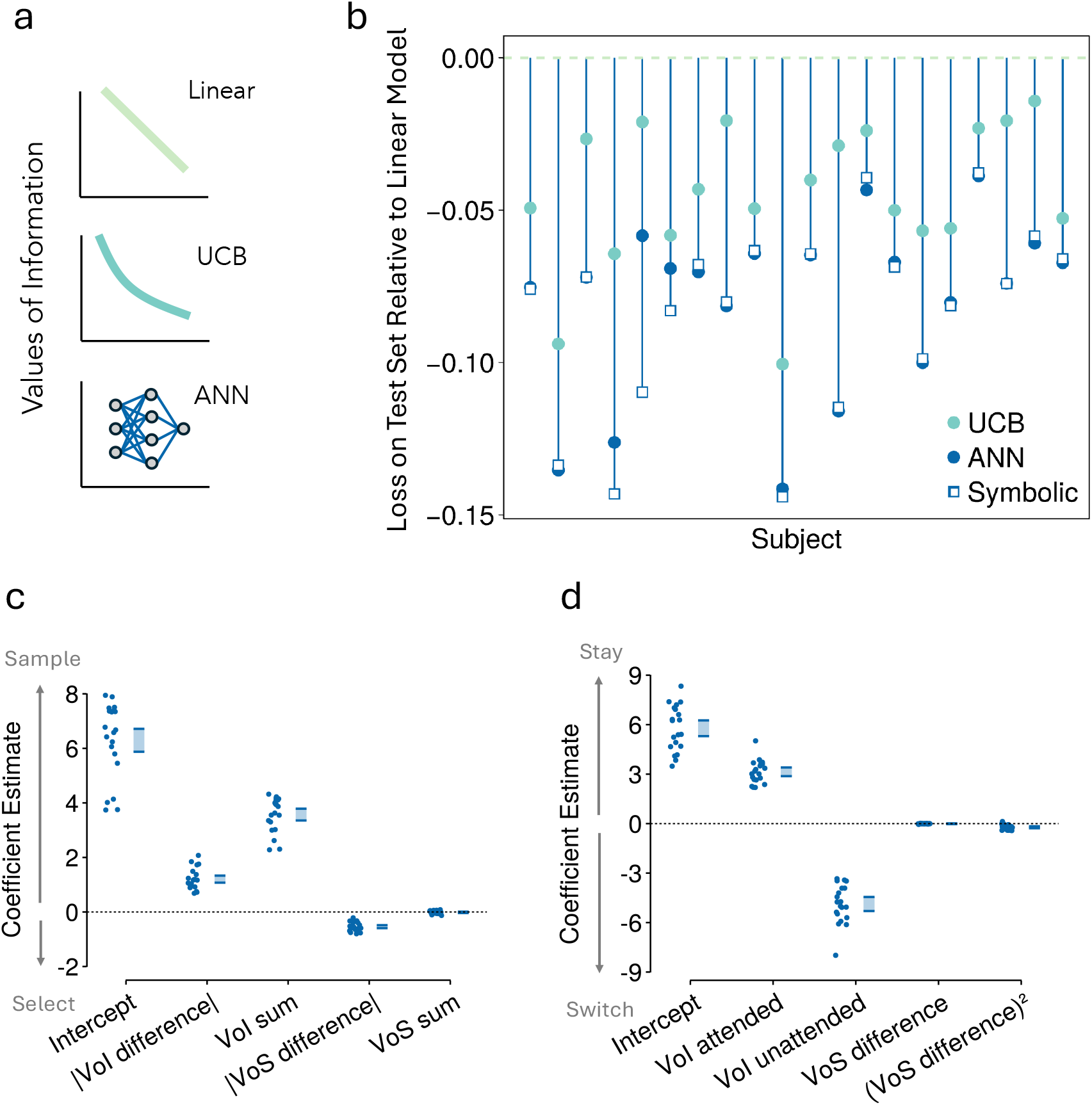
Model comparison and behavioral predictions. a, Three approaches to computing the value of information: linear function (top), Upper Confidence Bound (UCB) algorithm (middle), and artificial neural network (ANN; bottom). b, Model comparison across participants showing loss relative to the linear model. Each vertical line represents a participant, with light blue circles (UCB model), dark blue circles (ANN model), and open squares (symbolic model derived through symbolic regression) showing the relative loss for each model type. Lower values on the y-axis indicate better model fit. c, Regression analysis predicting the probability of sampling versus selecting. Each dot represents a subject-specific random effect estimate from a mixed-effects logistic regression model, with blue rectangles showing the ±95% confidence intervals of the fixed effects. Predictors include the intercept, absolute difference in value of information (|VoI difference|), sum of value of information (VoI sum), absolute difference in value of selection (|VoS difference|), and sum of value of selection (VoS sum).d, Regression analysis predicting the probability of staying versus switching. Subject-specific random effect estimates (dots) and fixed effect ±95% confidence intervals (blue rectangles) for predictors including intercept, value of information for attended patch (VoI attended), value of information for unattended patch (VoI unattended), and value of selection differences.

Once the ANN estimates the value of information for sampling each patch, the model computes the overall value for four potential actions (Fig. 2c, right). For the two sampling actions, the value of staying to sample the currently attended patch is its ANN-estimated value of information. The value of switching to sample the unattended patch is its ANN-estimated value of information, reduced by a fitted cost of switching. For the two selection actions, the model compares the current subjective estimates of red dots (*ρ*) in the attended and unattended patches. The value of selecting the attended patch is based on the difference in these subjective estimates (*ρ*_*attended*_ − *ρ*_*unattended*_), and correspondingly for selecting the unattended patch (*ρ*_*unattended*_ − *ρ*_*attended*_).

These four action values are then scaled by a temperature parameter and transformed via a softmax function to yield the probability of choosing each action.

To assess whether our hybrid modeling approach could accurately recover known, arbitrary value-of-information functions, we used a simulation-recovery procedure. We simulated data from four different agents, each using a distinct non-linear function to compute their value of information. For each simulated agent, we then applied our hybrid modeling approach to recover the underlying function from their behavior. The recovered functions showed very high correlations (r ¿ 0.95) with the true generating functions across different sample sizes, from 4,000 to 12,000 observations (Supplementary Fig. 1). This validation demonstrates that our hybrid approach can reliably recover complex value-of-information computations from behavioral data, giving us confidence in applying this method to understand human behavior.

We then performed feature selection to identify which task variables were essential for the ANN to compute the value of information. We initially considered a set of candidate input features for the ANN, including the current gaze position (i.e., which patch was attended), whether it was the first visit (i.e. a dummy variable that remains true as long as the participant continues to sample the patch they first attended at the start of the trial and becomes false upon the first switch to the other patch within that trial), the number of pieces of evidence (N) for both the attended and unattended patch (noting that for an unvisited patch, its N is formed by its initially visible green dots, while for a visited patch, N indicates the number of both red and black dots. In all cases N indicates the number of non-grey dots), the subjective proportion of red dots (*ρ*) for both patches, and the number of green dots in the blocked option. Using cross-validation, we systematically assessed how model performance changed when individual features were removed from the input vector. This analysis revealed that four features were critical: the current gaze position, whether it was the first visit, and the number of pieces of evidence (N) in the attended and unattended patch. Removing any of these features led to significant drops in model performance (Supplementary Fig. 2; see Methods for details).

Having validated our method and identified the input features, we next compared how well each approach to computing the value of information (e.g. linear, UCB, or hybrid-ANN) could account for participants’ behavior. Using a cross-validation approach, we found that for each participant the hybrid model achieved a better fit than the other two models for all participants (Fig. 3b). We also considered a Symbolic Model in this model comparison (Fig 3b), which we detail in the next section. This suggests that the ANN was able to find an alternative strategy that the first two competing models did not capture.

We then assessed how the ANN-derived value of information affected sampling decisions (Fig. 3c). Firstly, we used a mixed-effect logistic regression model to predict the probability of sampling information versus selecting a patch. In other words, we estimated the probability that participants would take either of the first two actions (Fig. 1a, bottom: staying to continue sampling from the current patch or switching to sample from the other patch) as opposed to either of the last two actions (Fig. 1a, bottom: selecting the currently attended option or the currently unattended option). We found that the probability of sampling was positively associated with the sum of the ANN-derived value of information of both options (*β* = 2.860, *s*.*e*. = 0.167, *z* = 17.16, *P* = 5.25 × 10^−66^), and with the unsigned difference in value of information between the two options (*β* = 1.292, *s*.*e*. = 0.085, *z* = 15.16, *P* = 6.14 × 10^−52^). This suggests that participants were more likely to sample information when more information was potentially obtainable in the near future, and when the discrepancy between the amount of information obtainable from the two options was large. From here on we refer to the value of information as being higher when these two factors were higher.

Secondly (Fig. 3d), we used a mixed-effect logistic regression model to predict the probability that one or other of the two different options would be sampled for information (Fig. 1a bottom, right versus left): staying – continuing to attend the currently attended patch – versus switching to attend to the alternative patch. We found that the probability of staying was positively associated with the value of information of the currently attended patch (*β* = 3.971, *s*.*e*. = 0.434, *z* = 9.14, *P* = 6.03 × 10^−20^) and negatively associated with the value of information of the unattended patch (*β* = −5.880, *s*.*e*. = 0.456, *z* = −12.89, *P* = 5.13 × 10^−38^), indicating that participants were more likely to sample from the currently attended patch when its estimated value of information was high relative to the unattended option.

Finally, we examined how participants made their final selection between the attended and unattended patches (Fig. 1a top; Supplementary Fig. 3). Using a mixed-effects logistic regression, we found that participants’ choices were strongly influenced by the proportion of red dots in both patches, with a positive effect for the proportion difference between attended and unattended patch (*β* = 4.572, *s*.*e*. = 0.336, *z* = 13.60, *P* = 3.77 × 10^−42^) and a positive effect for the proportion sum (*β* = 0.631, *s*.*e*. = 0.104, *z* = 6.08, *P* = 1.19 × 10^−9^). The value of information also influenced these selection decisions: participants were less likely to select the attended patch when its value of information was higher than the value of information of the unattended patch (*β* = −0.782, *s*.*e*. = 0.159, *z* = −4.91, *P* = 9.16 × 10^−7^). This pattern suggests that participants interpreted high remaining information value as uncertainty about the true proportion of red dots, making them less likely to select patches with high remaining uncertainty. It is also consistent with an account of choice in which participants seek to minimize uncertainty about the option that they will ultimately choose ^37^.

In summary, cross-validation analyses revealed that, for each participant, the hybrid-ANN model provided a superior account of information sampling behavior compared to both linear and UCB models (Fig. 3b). Subsequent regression analyses demonstrated how this ANN-derived value of information guided participants’ decisions (Fig. 3c-d): it significantly predicted when participants chose to sample further information versus make a final selection, and where they directed their sampling (i.e., whether to stay with the current patch or switch to the alternative). While final selection decisions were primarily driven by the perceived proportion of red dots, the value of information also played a secondary role, with participants tending to avoid options with a high remaining value of information (implying uncertainty about the number of red dots associated with the option; Supplementary Fig. 3).

### The ANN integrates evidence from both patches to compute value of information

Our behavioral analyses demonstrate that the hybrid-ANN approach has a better predictive performance than the linear and UCB models. To gain insight into the nature of the computation learned by the ANN, we first performed a qualitative analysis of the ANN-learned function (Fig. 4).

**Fig. 4.**
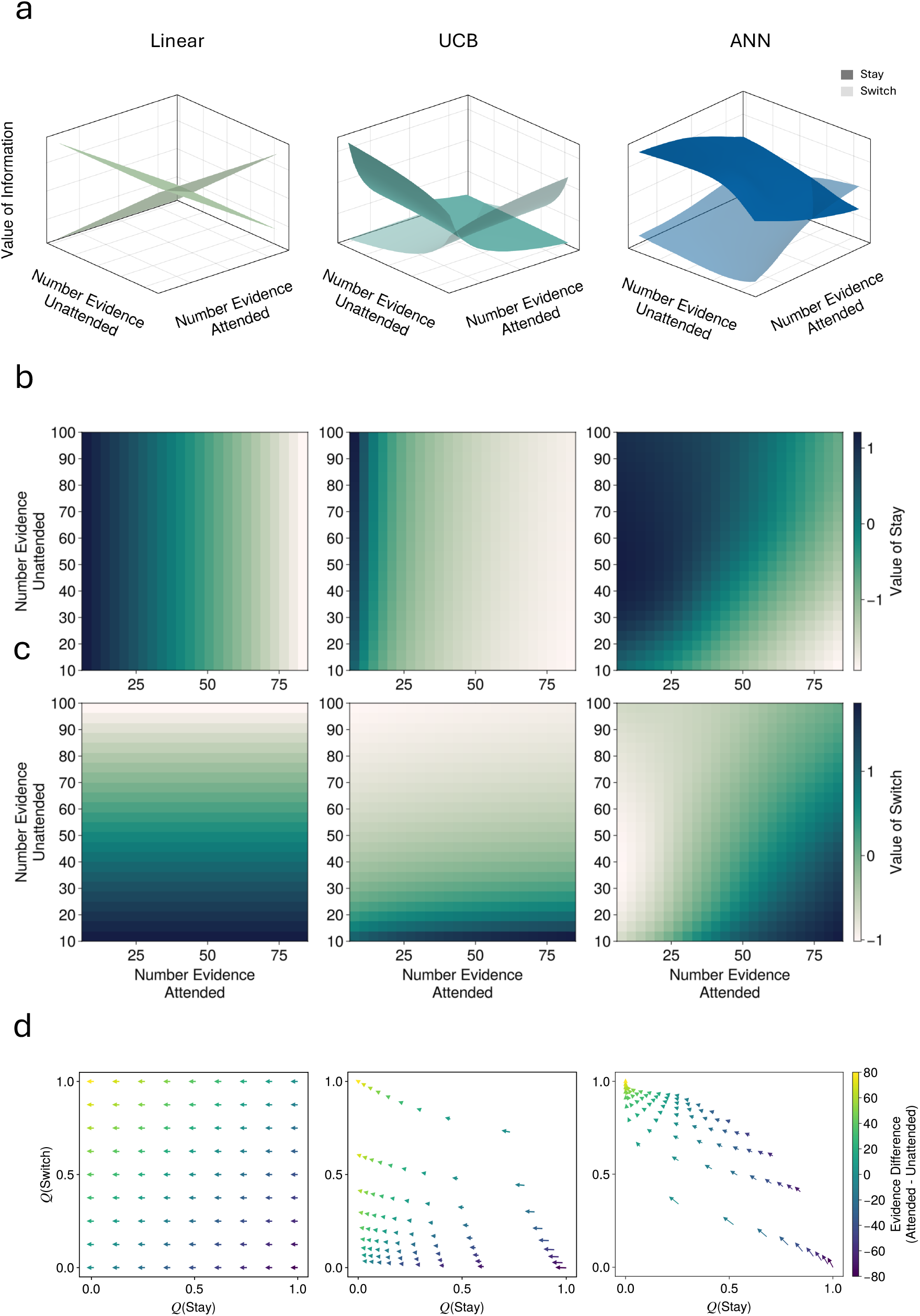
Visualization of value of information functions across computational models. a, Three-dimensional surface plots showing how the value of information varies with the number of revealed dots in both the attended and unattended patches. Each model (Linear, UCB, and ANN) is represented by two surfaces: darker color for the value of staying with the attended patch and lighter color for the value of switching to the unattended patch. b, Heatmaps showing the value of staying as a function of evidence collected from both patches. Color intensity represents the magnitude of the value, with the scale shown on the right. c, Heatmaps showing the value of switching as a function of evidence collected from both patches. d, Vector field plots showing how collecting an additional sample from the attended option affects both the value of staying (x-axis) and switching (y-axis). Each arrow represents the transition from current values (arrow origin) to updated values (arrow tip) after collecting one new sample. The color of the arrows indicates the difference in evidence between attended and unattended patches according to the color scale on the right, while the length of the arrow indicates the size of the update.

We started by simply visualizing the input-output relationship of the three functions (linear, UCB, and ANN) as a surface in 3D space relating the number of pieces of evidence (N) available in the attended option and in the unattended option to the value of information associated with staying and continuing to sample the attended option, and to the value of information associated with switching to the alternative unattended option (Fig. 4a). It should be noted that, because the participants might take another course of action, actually selecting an option, the value of staying to gather more information about the current option and the value of switching to gather information about the alternative option, are not simply inverses of one another. We then created heatmaps for each model showing the value of gathering information from the attended (value of staying; Fig. 4b) and unattended patches (Fig. 4C; value of switching) as a function of the evidence available to both the attended and unattended patches. Figure 4B shows distinct patterns in how the three models integrate information from both patches. The linear model’s gradient varies exclusively along one dimension, the attended evidence, when computing the value of staying, and the unattended evidence when computing the value of switching. The UCB model shows a similar predominant dependence on one dimension, though with some modulation by the second dimension. This behavior is an expected consequence of the models’ core assumptions. The linear model calculates the value of information for each patch in isolation, based solely on the evidence accumulated for that specific patch. Therefore, the decision to stay or switch is influenced only by the state of the patch being considered for the next sample. In contrast, UCB evaluates the informational value of an option in relation to the overall information gathered from all options. This means that the evidence from both patches contributes, albeit to different degrees, when assessing the value of sampling either the attended or the unattended option. The ANN-derived value of stay, shows clear dependencies on both dimensions: when computing the value of staying, it increases with attended evidence but simultaneously decreases with unattended evidence, resulting in a diagonal gradient pattern. This pattern indicates that the ANN, unlike the Linear model, learns to integrate evidence from both patches. While the UCB model also exhibits a two-dimensional influence, the ANN assigns a more substantial role to unattended evidence when computing the value of staying (contributing to the observed diagonal gradient).

To quantify this pattern, we performed a linear regression analysis to estimate how strongly each model’s output depends on attended versus unattended evidence (Supplementary Fig. 4a). For the value of stay computation, the Llinear model shows an exclusive dependence on attended evidence (*β*_*attended*_ = 1.0) with no influence of unattended evidence (*β*_*unattended*_ = 0.0). The UCB model shows a similar but less extreme pattern, with a strong weight on attended evidence (*β*_*attended*_ = 0.8) and a modest one on the unattended evidence (*β*_*unattended*_ = 0.2). The ANN, in contrast, shows more balanced weights for both attended (*β*_*attended*_ = 0.7) and unattended (*β*_*unattended*_ = 0.6), confirming our visual observation that it integrates information from both patches more evenly when computing the value of staying. A similar pattern emerges for the value of switch computation, where the ANN again shows more balanced weighting of evidence from both patches compared to the Linear and UCB models (Fig. 4c; Supplementary Fig. 4b).

To further analyze the difference between the ANN and the other two models, we visualized in Figure 4d how collecting an additional sample from the attended option affects both the value of staying and switching. Each arrow in the plot represents the transition from current values (arrow origin) to updated values (arrow tip) after collecting one new sample. Note that the value of staying and the value of switching are computed separately rather than being inverses of each other, because both are weighed against the value of selecting either patch in the final softmax comparison across all four possible actions. In the Linear and UCB models, these arrows are predominantly horizontal, indicating that new samples from the attended option strongly affect the value of staying but have minimal impact on the value of switching. In contrast, the ANN model has more diagonal arrows, revealing that new information from the attended option influences both the value of staying and switching. Specifically, the diagonal pattern suggests that as evidence accumulates from the attended patch, the value of staying decreases while the value of switching increases by a similar amount. This coordinated change means that while there may be relatively little change in the overall probability of continuing to sample (versus making a final selection), there is an increase in the probability of switching attention to sample the unattended patch. Overall, these results suggest that the ANN has learned a function that uses information from both the attended and unattended patches to guide sampling decisions.

### The ANN can be transformed into an interpretable symbolic function

While the qualitative analyses reported so far provide valuable insights into how the ANN operates, they do not provide a precise mathematical description of the computation it performs. To obtain a quantitative interpretation of the ANN’s learned function, we turned to symbolic regression, a method that can discover interpretable mathematical expressions that approximate complex nonlinear functions ^38^. This approach is particularly powerful for neural network interpretation for two reasons. First, it can find human-readable expressions that capture the essential computations performed by the network while eliminating the complex transformations that typically obscure these computations in the network’s internal representations^39^. Second, it can dramatically reduce the number of parameters (in this case, 7592 trainable units in the ANN) to a few, interpretable features.

We extracted a mathematical expression for the value of stay and one for the value of switch. For the value of stay, we extracted the following expression 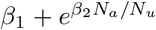 (Fig. 5a), where *N*_*a*_ and *N*_*u*_ represent the number of dots revealed in the attended and unattended patches respectively. For the value of switch, we found 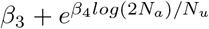 (Fig. 5a). As can be seen from these equations, *β*_1_ acts as a global offset for the value of stay, effectively controlling the balance between undirected exploration and maintaining attentional focus on the currently attended option (see Fig. 5c, left). *β*_2_ modulates how strongly the value of stay is influenced by the interaction between attended and unattended evidence (Fig. 5c, second from left). Similarly, *β*_3_ acts as an intercept for value of switch and *β*_4_ determines the sensitivity of the value of switch computation to the relative amounts of evidence between patches (Fig. 5c, right). Notably, while these equations contain four core parameters, our feature selection analysis revealed that the visit number (first visit of the patch vs subsequent visits) was critical for optimal ANN performance. Consequently, each parameter is fitted independently for first visits versus subsequent visits, yielding eight total parameters that capture distinct sampling strategies across different phases of exploration. These interpretable parameters provide insight into individual differences in sampling strategies (Supplementary Fig. 7c).

**Fig. 5.**
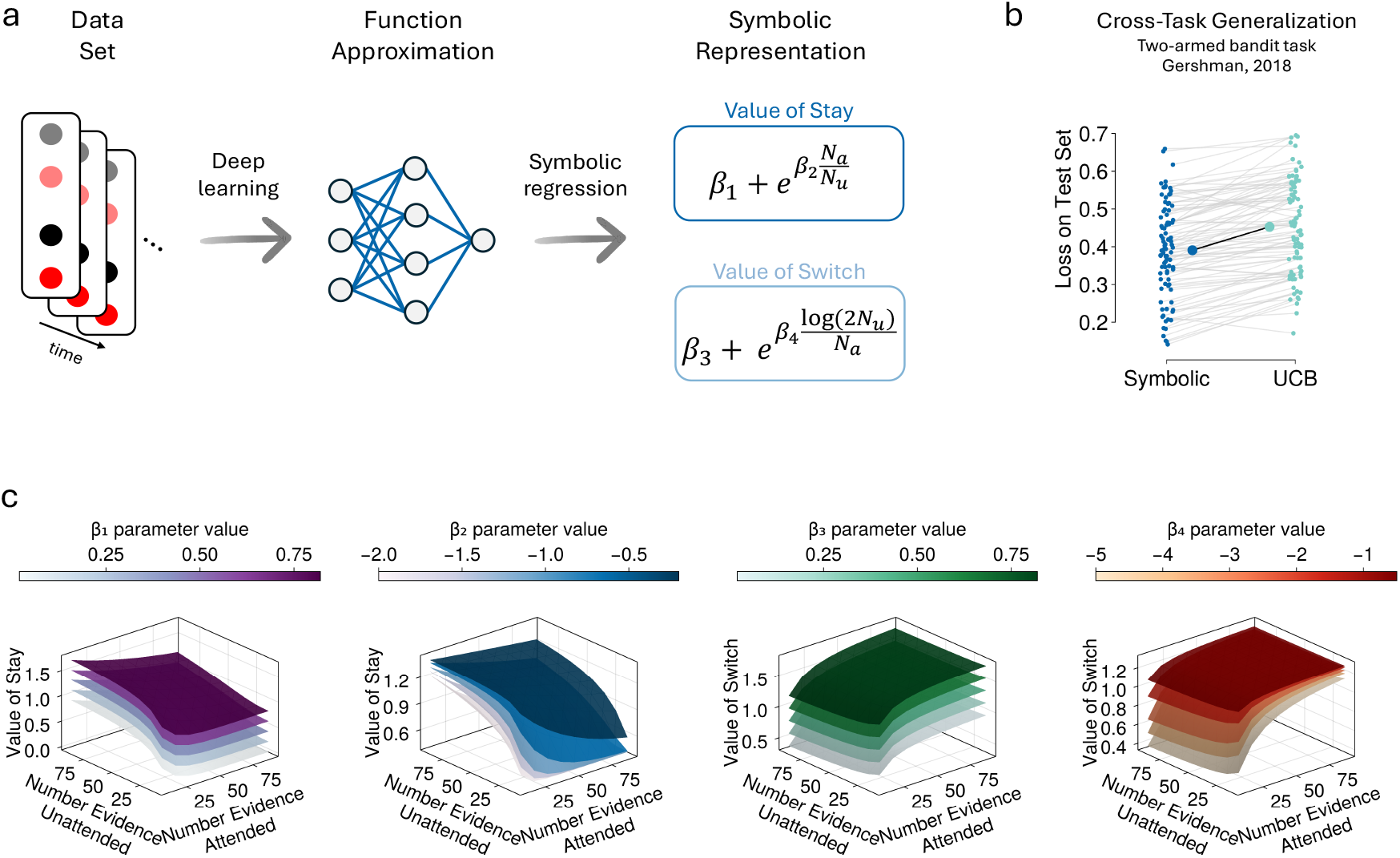
Symbolic representation of the ANN-derived value of information function. a, Process of deriving interpretable mathematical expressions from behavioral data. Left: Schematic of the dataset containing patterns of dots across time. Middle: Function approximation using a deep neural network. Right: Symbolic representation showing the mathematical equations derived through symbolic regression for the value of staying and the value of switching, where *N*_*a*_ and *N*_*u*_ represent the number of dots in the attended and unattended patches, respectively. b, Cross-task generalization comparison. Scatter plot showing loss on a test set from a two-armed bandit task (Gershman, 2018) for both the symbolic model (left) and UCB model (right). Each gray line connects performance of both models for the same participant. c, Parameter sensitivity analysis. Three-dimensional surface plots showing how the value functions change with different parameter values. Each surface represents the function with a specific parameter value according to the color scales above each plot.

To examine whether these symbolic functions accurately capture the function learned by the ANN, we replaced the ANN component in our hybrid model (Fig. 3a, bottom) with these newly discovered functions and assessed how well this new variant of the model explained participant behavior using the approach. We found the symbolic-hybrid model and ANN-hybrid had comparable power to explain participant behavior (Fig. 3b), suggesting we had successfully discovered a transparent and interpretable mathematical relationship between evidence and the value of information.

Finally (Fig. 5b), we tested whether these symbolic functions capture general principles of information sampling rather than task-specific features. We evaluated performance of the symbolic model on an independent dataset where participants completed a two-armed bandit task ^19^. Participants in this study performed a very different task from ours (e.g. they repeatedly chose between two options and received point rewards), yet in both tasks participants had to strike a balance between exploration and exploitation to maximize their rewards. Remarkably, our symbolic functions outperformed the UCB model in predicting participants’ choices also in this different context (Wilcoxon signed-rank test: *W* = 15.0, *n* = 89, median difference = −0.07, *P* = 4.31 × 10^−16^). We noticed that one simple difference between the standard UCB formulation and our symbolic model is that the UCB lacks an offset parameter while our symbolic model includes one. To ensure that this superior performance was not simply due to the model’s ability to adjust the baseline offset, but rather reflected a fundamental difference in the shape of the value-of-information function, we tested an UCB model that included an offset parameter in its exploration bonus computation. Even when compared to this more flexible UCB variant, the symbolic model still demonstrated significantly better predictive performance (Wilcoxon signed-rank test: *W* = 286.0, *n* = 89, median difference = −0.03, *P* = 2.21 × 10^−12^). These results suggest that the equations discovered with symbolic regression capture fundamental aspects of human exploration behavior that are not specific to the information sampling task we used and cannot be accounted for by simple parametric extensions to existing models.

### The ANN-derived value of information can predict neural activity

Our behavioral analyses demonstrate that the ANN-derived value of information better predicts participants’ behavior (Fig. 3b). Next, we sought to understand whether this value of information could also predict neural activity. Participants performed the behavioral task while undergoing ultra-high field fMRI recordings of the blood oxygen level dependent (BOLD) signal. To achieve high spatial resolution (1mm isotropic voxels), our functional imaging protocol used a limited field of view that captured key regions of interest in the midbrain, brainstem, and interconnected cortical areas (Supplementary Fig. 6), rather than whole-brain coverage. We used these recordings to investigate brain activity that represented the main task variables. We employed a general linear model (GLM) analysis across the whole brain volume scanned. The GLM used the value of information as a predictor of neural activity, while controlling for other task-relevant variables such as outcome, and value of selection. We compared the goodness of fit when using three different value-of-information computations: our ANN-derived estimates, the linear model estimates, and the UCB model estimates. For each voxel within our imaging field of view, this GLM analysis yielded a predicted BOLD timeseries for each of the three VOI models. We then calculated the discrepancy (Mean Squared Error, MSE) between the observed BOLD timeseries and the model-predicted timeseries on a per-voxel basis. This provided a quantitative measure of each model’s ability to fit the neural data in each voxel. Note that (unlike for behavior in Fig. 3b), this model comparison fits the same number of parameters per model when predicting the BOLD signal: it takes the model output value of information (holding the number of internal parameters in the model fixed) and uses them as parametric modulators in a conventional GLM analysis of fMRI data. Consistent with our behavioral findings, the ANN-derived value of information demonstrated a better overall fit to the neural data across the brain. Specifically, when comparing the distribution of these per-voxel MSEs, the ANN-derived VOI resulted in significantly lower MSEs than both the linear and UCB models (Wilcoxon signed-rank test: median *MSE*_*ann*_–*MSE*_*linear*_ = −82.42, *P* = 2.15 × 10^−12^; median *MSE*_*ann*_–*MSE*_*ucb*_ = −90.77, *P* = 5.74 × 10^−13^; Supplementary Fig. 5). This suggests that the computational principles captured by our hybrid ANN model not only better describe participants’ behavior but also more accurately reflect the underlying neural computations driving information sampling decisions. We then investigated activity in specific brain regions.

### Anterior Insula and Anterior Cingulate Cortex covary with the ANN-derived Value of Information

To identify the neural correlates of the ANN-derived value of information, we conducted two complementary analyses. First, we performed a whole-brain GLM analysis to explore cortical regions most strongly associated with the ANN-derived value of information. Second, we conducted targeted analyses on pre-defined regions of interest (ROIs) at the origins of the neuromodulatory systems, DRN, LC, VSN, VTA, and SN (Fig.1D). The neuromodulators associated with each these nuclei have been proposed previously as mediators of the impact of uncertainty on decision-making ^3,20–26^. In general, unlike the cortical regions, they are too small to survive cluster correction analysis strategies that incorporate a thresholding criterion based on the spatial extent of the activity ^27,40,41^.

The whole-brain analysis revealed that activity in the anterior insula (AI) and anterior cingulate cortex (ACC) was associated with the ANN-derived value of information (Fig. 6a and Fig. 6b). AI activity exhibited a negative association with the difference between value of stay and value of switch (Fig. 6a top) and a negative association with the sum of value of stay and value of switch (Fig. 6b top); in other words, AI activity increased as the value of switching attention increased and decreased as the value of maintaining attention at the current location increased. The ACC had the same pattern (Fig. 6b). These computational signatures appear crucial for guiding participants’ decisions about whether to stay with the current patch or switch to the alternative (Fig. 1b, bottom), and whether to keep sampling information or stop the learning process and initiate an option selection (Fig. 1a). To test whether the superior model fit extended to these two specific regions, we compared the MSE between models within AI and ACC ROIs. Consistent with the whole-brain results, the ANN-derived VOI showed significantly lower MSE than both linear and UCB models in both regions (AI: ANN vs Linear MSE difference = −175.183, *P* = 1.94 × 10^−12^; ANN vs UCB MSE difference = −200.76, *P* = 1.01 × 10^−12^; ACC: ANN vs Linear MSE difference = −57.088, *P* = 7.02 × 10^−5^; ANN vs UCB MSE difference = −50.246, *P* = 1.42 × 10^−5^).

**Fig. 6.**
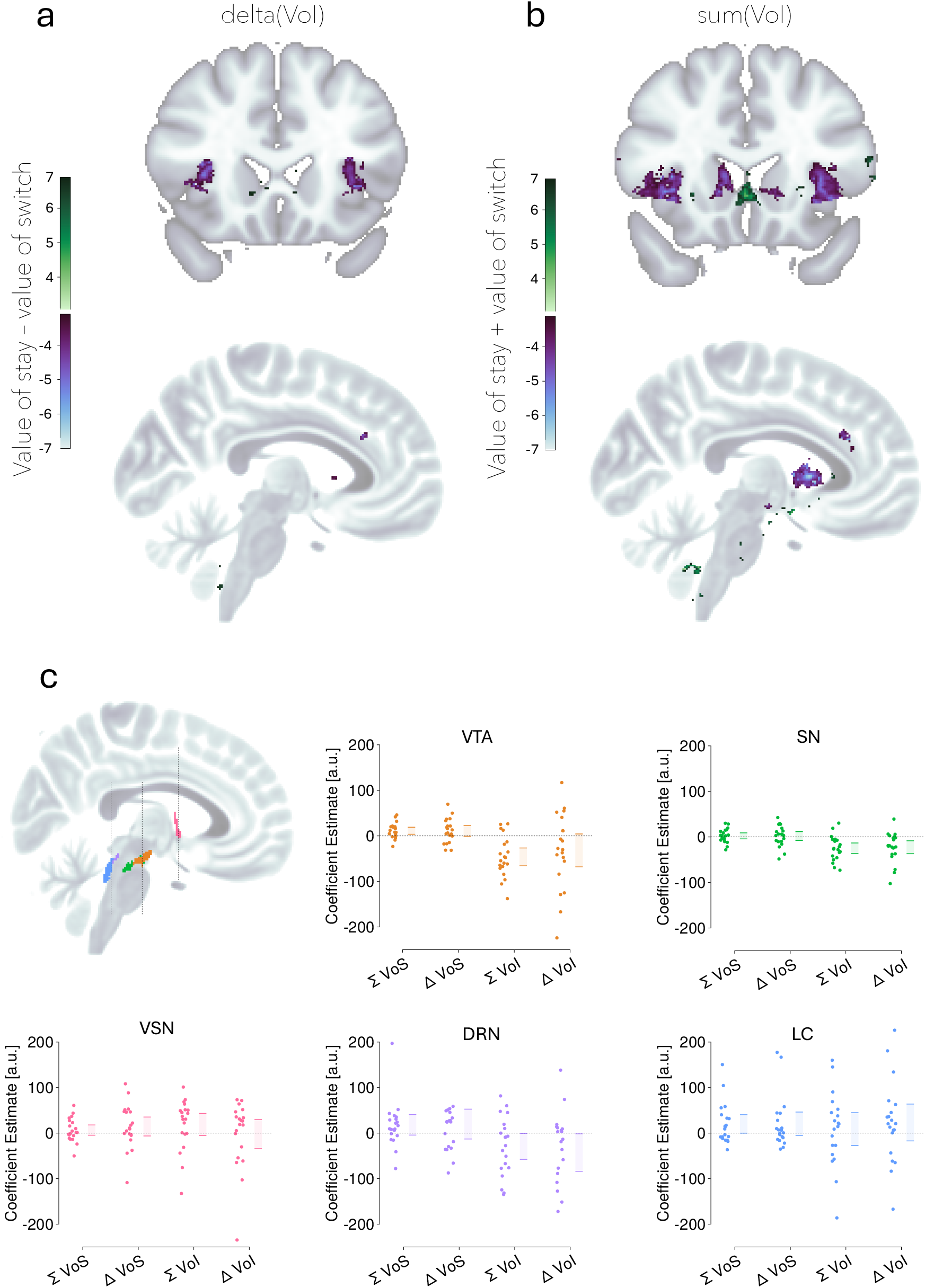
Neural correlates of ANN-derived value signals. a, Whole-brain analysis showing regions where BOLD activity correlates with the difference between the value of staying and the value of switching (VoI Stay - VoI Switch). b, Whole-brain analysis showing regions correlating with the sum of the value of staying and switching (VoI Stay + VoI Switch). Color bars indicate Z-statistic values; thresholded at *Z >* 3.1, cluster-corrected *P <* 0.001. c, ROI analysis results for neuromodulatory nuclei. Left: Sagittal view showing locations of VTA (orange), SN (green), VSN (pink), DRN (purple), and LC (blue). Right: Coefficient estimates (effect sizes from weighted mixed-effects models on beta values) for regressors representing the sum (Σ) and difference (Δ) of the value of selection (VoS) and value of information (VoI) within each ROI. Each small dot represents an individual participant’s random effect estimate plus the fixed effect; shaded bars indicate the 95% confidence interval of the fixed effect (group mean).

Analysis of the subcortical ROIs revealed a distinct pattern in the VTA (Fig. 6c), which showed positive coding of the sum of value of selecting the attended and unattended patch (*β* = 11.295, *s*.*e*. = 3.929, *z* = 2.87, *P* = 0.020) but negative coding for the sum of value of information (*β* = −46.111, *s*.*e*. = 9.978, *z* = −4.620, *P* = 1.93 × 10^−5^). This activity pattern would be sufficient to guide participants’ decisions about whether to continue sampling information from the patch they are attending or to make a final selection (Fig.1a, bottom right versus top right).

The SN exhibited a pattern similar to that observed in the cortical regions AI and ACC (Fig. 6c). Activity in the SN was negatively associated with both the sum of the value of information (*β* = −24.863, *s*.*e*. = 5.924, *z* = −4.19, *P* = 6.38 × 10^−5^) and the difference in the value of information between staying and switching (*β* = −22.665, *s*.*e*. = 7.243, *z* = −3.129, *P* = 0.008). This suggests the SN, like AI and ACC, tracks aspects related to the overall potential for information gain and the relative informational value of the current versus alternative option.

There has been particular interest in the possibility that the LC, and the noradrenergic system with which it is linked, the DRN, and its serotonergic system, or the VSN, associated with the cholinergic system, encode uncertainty. Our high-field fMRI recordings gave us a unique opportunity to test these hypotheses in humans. We first examined whether LC, DRN, or VSN activity can predict the ANN-derived value of information, an index closely, but inversely, related to uncertainty.

We further examined activity related to the sum and difference of the value of information and the sum and difference of the value of selection within the anatomically defined VSN, DRN, and LC ROIs (Fig. 6c). In contrast to VTA and SN, we found no significant association between BOLD activity in VSN, DRN, or LC and any of these four regressors. All p-values in the ROI analyses have been Bonferroni corrected for multiple comparisons across ROIs. These results suggest that, within the sensitivity limits of our measurement and analysis approach, these specific neuromodulatory nuclei do not strongly encode these particular decision variables related to the value of information or selection in this task.

To comprehensively assess model performance across subcortical regions, we compared the MSE between models within all five subcortical ROIs (VTA, SN, DRN, LC, VSN). First-level GLMs fitted using ANN-derived VOI demonstrated significantly lower MSE than those fitted using linear-derived VOI across all subcortical regions, and significantly lower MSE than those fitted using UCB-derived VOI in four of the five regions (all except LC after Bonferroni correction; Table S1. This consistent superiority of the ANN-derived approach across diverse neuromodulatory nuclei further supports the conclusion that the computational principles captured by our hybrid model better reflect the underlying neural mechanisms of information valuation, even in regions where the VOI signal itself may not reach statistical significance.

### AI and ACC Activity Reflects Information Value and Predicts Sampling Behavior

Our preceding analyses established that the ANN-derived value of information (VOI) predicts both sampling behavior (Fig. 3b) and BOLD activity (Supplementary Fig. 5). This implies a crucial, though as yet untested, link: the trial-by-trial variations in neural activity within VOI-encoding regions should directly correspond to trial-by-trial variations in sampling behavior. To explicitly test this implication and to investigate whether the representational structure of activity in these regions—beyond average activation levels—relates to behavior, we employed Representational Similarity Analysis (RSA). We reasoned that if a brain region computes decision variables guiding information sampling, then the similarity of its neural activity patterns across pairs of trials should mirror the similarity in the extent of sampling behavior observed on those same trials (Fig. 7a). Specifically, based on our univariate results (Fig. 6a and 6b), we hypothesized that the neural patterns in AI and ACC would exhibit this correspondence with sampling duration.

**Fig. 7.**
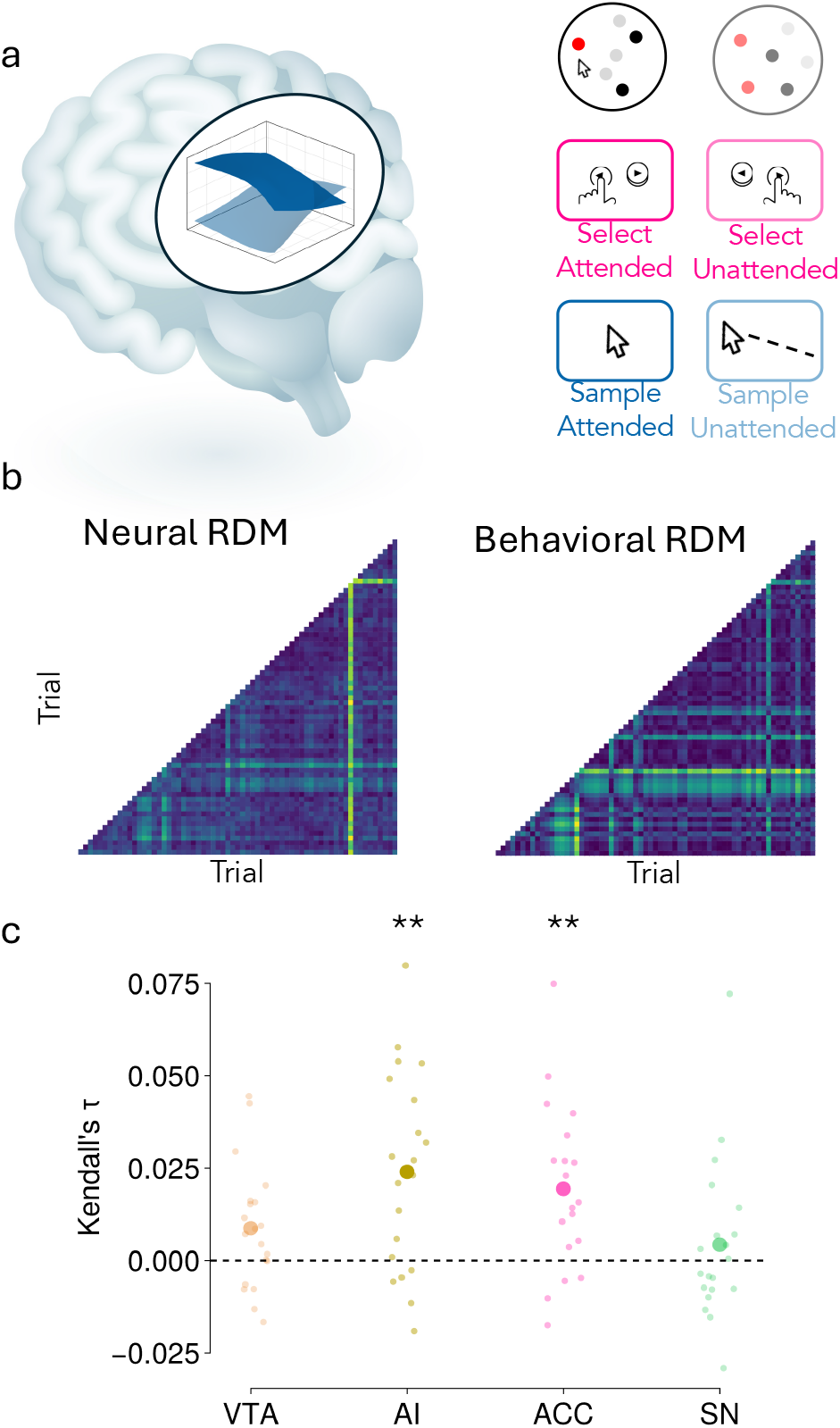
Representational Similarity Analysis links neural patterns to sampling behavior. a, Schematic illustrating the relationship between brain activity and behavior. b, Example Neural RDM (left, based on multi-voxel BOLD activity patterns) and Behavioral RDM (right, based on the number of samples taken per trial) for a participant session. Each cell represents the dissimilarity between a pair of trials; higher values (yellow) indicate greater dissimilarity. c, Kendall’s *τ* correlation coefficients quantifying the relationship between neural and behavioral RDMs for VTA, AI, ACC, and SN. Neural RDMs for AI, ACC, and SN were derived using masks matched in shape and size to the VTA ROI to control for region size. Each small dot represents the correlation calculated for one participant session; larger dots indicate the mean correlation across participants. Asterisks denote significant positive correlations across participants (Wilcoxon Signed Rank Test, \*\**P <* 0.01, Bonferroni corrected).

To test this, we constructed Representational Dissimilarity Matrices (RDMs) for each participant and session (Fig. 7b). A behavioral RDM captured the dissimilarity in sampling duration (number of samples taken) between pairs of trials using Euclidean distance. Correspondingly, a neural RDM captured the dissimilarity in BOLD activity patterns between the same pairs of trials, also using Euclidean distance. We focused on the four regions implicated in our univariate analyses (VTA, SN, AI, ACC), and to control for potential confounds related to the size and shape of regions of interest, we used masks matched to the VTA’s shape, positioned at the center of each respective target region.

We then calculated the Kendall’s *τ* correlation between the flattened neural and behavioral RDMs for each session. Statistical testing across participants revealed significant positive correlations between neural pattern similarity and behavioral similarity in both the AI (Signed Rank Test, *τ* = 0.024, *P* = 0.0006, Bonferroni corrected *P* = 0.0024) and in the ACC (*τ* = 0.019, *P* = 0.0004, Bonferroni corrected *P* = 0.0017), but not in the VTA (*τ* = 0.008, *P* = 0.072) or SN (*τ* = 0.004, *P* = 0.46, Bonferroni corrected *P* = 1.0) (Fig. 7c). This suggests that the patterns of activity within AI and ACC, beyond their average activation levels, contain information related to the amount of information participants choose to sample on a given trial. The lack of significant correlation in VTA and SN, despite their univariate associations with decision variables, indicates their trial-by-trial activity patterns may relate less directly to the specific duration of sampling compared to AI and ACC.

## Discussion

Information seeking has been proposed to be a fundamental behavior in humans and other animals ^3,20,42,43^. Understanding how and why information is valued and sought can not only help us understand how people interact with information sources as variable as other individuals or the internet ^44,45^ but how information seeking can become pathological, maladaptive, or how it can affect mental wellbeing ^46,47^. At the same time, understanding the value of information can help us understand how people and animals resolve decisions; inherent within decision making is a trade-off between, on the one-hand information seeking in order to ensure that the best option is identified and selected and, on the other hand, quick and effective decision making that allows the individual to move on to the next behavior without forgoing other opportunities. If the decision maker deliberates for too long over the first decision, she may forgo the possibility of making a decision about subsequent opportunities. That the utility of information diminishes with time during decision making is consistent with observations of neural activity in the lateral intraparietal (LIP) cortex and superior colliculus that selects the target for an eye movement and brings about the end of deliberation and initiation of the movement ^1,4,9,10^. In the current study, however, we examined how the value of information is itself determined.

Four key parameters (Fig. 5a, c) determine value of information obtained from the current focus of attention as opposed to value of information at an alternative location. The first and third, *β*_1_ and *β*_3_, set the balance between maintaining attention at the current focus as opposed to exploring anywhere else. The second and fourth, *β*_2_ and *β*_4_, determine how the relative amount of information at the currently attended location and at a specific alternative influence maintenance of attention at the current location and of switching attention to the alternative location respectively. The characterization of value of information in the current study captures features of potential choices that are akin to the uncertainty that a decision maker might have about their value. However, the precise characterization of value of information was a result of the ANN approach that was adopted; the flexible ANN approach was, first, able to capture non-linear relationships between choice features and value of information and, second, it was able to discover the relationship in the absence of a prior hypothesis about the precise nature of the relationship. Critically, however, the nature of the relationship between the choice features and value of information did not remain implicit and uninterpretable within the ANN but, instead it was rendered interpretable by symbolic regression. While the limit to the range of situations in which the current model of value of information applies is yet to be determined, it is clear that the model has some generalizability; it was able to predict behavior in a very different task but which was also characterized by an explore-exploit dilemma ^19^ (Fig.5B).

There is a growing recognition that purely theory-driven models may sometimes oversimplify complex cognitive processes, while purely data-driven approaches like deep learning often suffer from a lack of interpretability. Recent work explores various ways to bridge this gap, often falling into two main streams. One stream focuses on leveraging neural networks while enhancing interpretability: this includes explicitly integrating ANNs within classic cognitive frameworks ^48^ or utilizing deep learning architectures specifically designed for interpretability or cognitive plausibility, such as tiny ^15^ or disentangled ^49^ recurrent neural networks. A second, less explored, stream aims for interpretability by discovering mathematical equations directly from behavioral data using equation discovery algorithms ^50^. The workflow we propose, integrates aspects of both these streams. First, we use a flexible ANN to learn complex input-output mappings without strong a priori constraints. Then, we use symbolic regression to distill these learned mappings into human-readable equations. A natural question that arises is why not apply symbolic regression directly to behavioral data rather than first fitting an ANN. There are several computational and methodological reasons for our two-stage approach. First, direct symbolic regression on the complex mapping from task state to behavior would involve an intractably large search space. By first learning this mapping with ANNs and then applying symbolic regression to the learned function, we factorize this complex problem into manageable components, dramatically reducing the search space from multiplicative to additive complexity. Second, our neural network architecture (Lipschitz-Bounded Deep Network ^51^) inherently enforces smoothness constraints on the learned function, providing stability that would not be guaranteed with direct symbolic regression. Third, and crucially, our symbolic regression operates within a broader cognitive model optimized via gradient descent. Embedding a non-differentiable genetic algorithm (symbolic regression) directly within this gradient-based optimization framework would create significant computational challenges. By replacing the value-of-information computation with a differentiable neural network, we transform this into a standard, tractable optimization problem. This approach thus combines the computational efficiency and stability of neural network training with the interpretability benefits of symbolic regression. This approach was central to the current study but it is likely to have wider applicability in cognitive science and neuroscience. We believe it holds significant potential for refining existing theories and discovering novel computational principles across diverse domains, from learning and decision-making to perception and social cognition.

As noted, value of information is related to uncertainty about a decision option and uncertainty has, in turn, been related, at one time or another, to all the major neuromodulatory systems ^3,20–26^. However, identifying where one system makes a specific or preeminent contribution remains difficult. By using ultra-high field 7T fMRI we recorded from the nuclei – VTA, SN, DRN, LC, and VSN – from which the neuromodulatory systems innervating the forebrain originate. By simultaneously recording from them all we hoped to ascertain if any had an especially strong relationship with value of information. This was the case for VTA (Fig.6C); the sum of the values of information of the options was associated with significant activity change. At the same, the sum of the values of the options – the number of red dots, which determined participants scores and final payouts – was also associated with significant activity modulation but in the opposite direction. Such a pattern of opposed activity change is consistent with VTA balancing the value of making a choice selection against the value of information to be gained from exploring the options further (arbitrating between the top and bottom rows of Fig.1A). The SN exhibited a related pattern of activity (Fig.6C); it also encoded the sum of the values of information of the options, but its activity was also determined by the difference in the values of information of the options. This suggests that SN encodes both the potential for overall information gain but also the relative value of information from the current focus of attention and an alternative (the bottom row of Fig.1A). Similar patterns were also found in two cortical regions, ACC and AI (Fig.6A, B). These are the two cortical regions known to project to, or adjacent to, VTA and SN as well as other midbrain and brainstem neuromodulatory nuclei and which have been reported to have activity that is related to that found in the neuromodulatory nuclei ^3,21,28–32,52,53^. There are also multisynaptic routes running between these cortical regions and VTA/SN via other brain structures such as the habenula ^54,55^. Importantly, multivariate analysis demonstrated that ACC and AI are especially intimately concerned with information evaluation; the multi-voxel pattern of activity in both ACC and AI covaried with the number of samples of information taken on each trial (Fig.7).

Previous studies have identified ACC and VTA and anatomical structures that interconnect them, such as habenula, in evaluation of information. Activity in individual neurons in both habenula and VTA reflects both the value of a stimulus in terms of the reward that it predicts ^56–58^ but also value of information ^20,21,26,59^ in line with the suggestion that VTA may compare the relative advantage to be gained from making a selection and seeking more information about a choice. Activity in ACC has also been linked to information seeking and initiation of behavioral change ^13,16,21,60–64^ consistent with the proposal that ACC might arbitrate when it is advantageous to seek more information about a current opportunity and when it is better to pursue an alternative. AI is relatively less investigated but, in line with the current findings, AI and activity in dopaminergic midbrain areas such as SN have been reported when people evaluate whether and when to initiate an action ^27,40^.

## Methods

### Subjects

Twenty participants (14 females), aged 19 to 32 years, completed the study. All participants were paid £15 per hour and an additional performance-dependent bonus of between £20 to £40 for rewards collected during the task. Each participant provided written informed consent at the beginning of each testing session. Ethical approval was given by the Oxford University Central University Research Ethics Committee (Ethics Approval Reference: R82877/RE001). Behavioral data from all participants were used for the analysis.

#### Experimental Task

On each trial, participants were presented with three patches of moving dots (100 dots per patch, radius = screen height/250, where screen height was scaled to 40% of the window height). Each dot’s true color was either red or black, but initially all dots were hidden under either green or grey covers. The green covers indicated dots whose colors would be revealed simultaneously upon the participant’s first visit to that patch, while grey covers indicated dots that would be revealed sequentially subsequently. The number of green-covered dots varied between patches and trials (between 70-95 dots), allowing separation of sampling time from patch uncertainty.

The sequence of each trial was as follows: (1) participants clicked a start button to initiate the trial (free response time); (2) they clicked on each patch to reveal the green-covered dots (free response time); (3) they then clicked a central sampling button to begin the information gathering phase (free response time); (4) in 20% of trials, one of the three patches was randomly blocked and could not be sampled or chosen; (5) participants could then freely sample information by hovering over patches, with grey-covered dots revealing their true colors sequentially every 150ms, for a maximum of 24 seconds when two alternatives were available or 33 seconds when all three patches were available; (6) after making their choice using an MRI-compatible button-box, participants received feedback through a “duel” animation (1300ms) followed by a win/lose image (1500ms). Each participant completed four 25-minute sessions and was incentivized to maximize correct choices with a potential bonus payment of £5 per session. The experiment was implemented using jsPsych (version 6.3.0) and custom plugins for stimulus presentation and response collection.

To make the task more engaging, we framed it as a medieval-themed game called “The Last Duel”, where participants could collect points by making correct choices. In this context, participants were told they lived in a hostile world where strangers might challenge them to duels. Their goal was to obtain the best weapon (sword *>* hammer *>* stone) to ensure victory in these duels, with their opponent always in possession of a hammer. The sword would guarantee victory (100% win rate), the hammer would give an equal chance (50% win rate), and the stone would always lead to defeat (0% win rate). The three patches represented weapon stores in town, and the proportion of red dots in each patch determined which weapon it contained: the patch with the highest proportion of red dots contained the sword, the intermediate proportion contained the hammer, and the lowest proportion contained the stone.

#### Behavioral Analysis

To analyze participants’ decision-making behavior, we employed mixed-effects logistic regression analyses to examine three key aspects of choice behavior. First, we investigated how participants decided between staying and sampling information at the current patch versus switching to sample information the other patch (Fig.1A, bottom). This analysis modelled the probability of staying versus switching as a function of the value of information for both the attended and unattended patches, while also accounting for the difference in means between patches and its quadratic term.

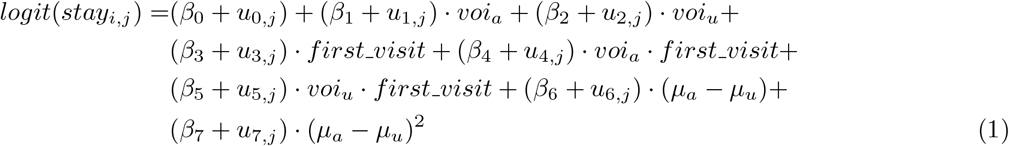

where *i* indexes observations, *j* indexes subjects, *β*_*k*_ are the fixed effects, and *u*_*k,j*_ are the subject-specific random effects. The random effects are assumed to follow a multivariate normal distribution: **u**_*j*_ ~ N (0, Σ), where Σ is the variance-covariance matrix of the random effects. *voi*_*a*_ and *voi*_*u*_ are the value of information for the attended and unattended patches, respectively. *µ*_*a*_ and *µ*_*u*_ are the observed proportions of red dots in the attended and unattended patches, respectively.

Second, we examined the factors influencing participants’ decisions between continuing to sample information versus making a final selection. This model included predictors capturing both the absolute difference and sum of the value of information between patches, including interactions with first visit, as well as the absolute difference and sum of the observed means.

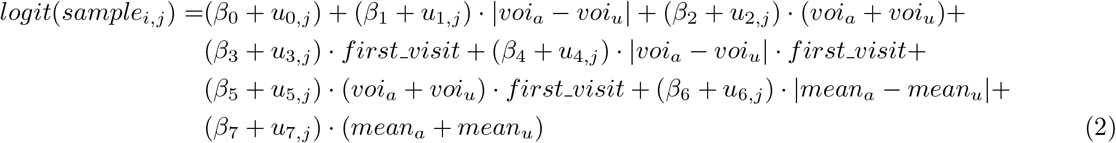

Third, we analysed participants’ final patch selections to understand what drove the choice between attended and unattended patches, using both the observed means and value of information of both patches as predictors. For each of these analyses, we compared three different approaches to computing the value of information: linear, upper confidence bound (UCB), and hybrid approaches. All models incorporated random effects for all predictors grouped by subject and used a logistic link function to account for the binary nature of the choices. This comprehensive modelling approach allowed us to dissect different aspects of the decision-making process while accounting for individual differences through the random effects structure.

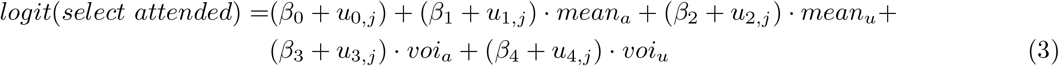

We fit each model for the three different approaches to computing the value of information. We used the library MixedModels.jl in Julia to fit the mixed-effects logistic regressions.

#### Optimal Model

We formulated the information sampling task as a Markov Decision Process (MDP) with states representing the agent’s knowledge about dots in each patch, actions corresponding to stay, switch, and selection decisions, and rewards capturing both sampling costs and final choice outcomes. Specifically, each state *s* contains: the current time step, the number of revealed red dots in each patch, the total number of revealed dots (red + black) in each patch, the current gaze position (left or right), and whether each patch has been visited. The action space consists of four possible actions: stay (continue sampling current patch), switch (move to the other patch), select attended patch, or select unattended patch.

The transition function *T* (*s*′|*s, a*) defines the probability of transitioning to the next state *s*′ when taking action *a* in the current state *s*. For sampling actions (stay/switch), the transition function follows a Beta-Binomial distribution where the probability of observing a new red dot is based on the current counts plus a uniform prior (*α* = 1, *β* = 1). Specifically, given *n* previously revealed dots (*r* red and *n* − *r* black) in the attended patch, the probability of observing a new red dot follows 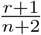, while the probability of observing a black dot follows 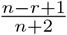. For selection actions, the episode terminates.

The reward function *R*(*s, a*) assigns different rewards based on the type of action. For sampling actions, the agent gets an immediate negative reward when she chooses sample an attritional piece of information from the option she was currently attending (costs for staying) and a bigger immediate negative reward when the agent chooses to switch, and sample from the other patch (cost of switch). Participants in the real task experienced a switch delay of 13.3 times longer than the stay delay (e.g. 2000ms versus 150ms). To reflect this difference in the reward function, we set the cost of switch to be 13 times bigger than the cost of stay). For selection actions (choosing either the attended or unattended patch), the reward is computed based on the probability that the selected patch has a higher true proportion of red dots. Specifically, given the current evidence for the attended patch (*α*_1_, *β*_1_) and unattended patch (*α*_2_, *β*_2_), we compute:

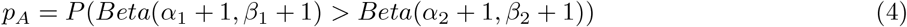

where the probability is calculated using a normal approximation based on the difference in means normalized by the square root of the sum of variances. The final reward for selecting the attended patch is then *p*_*A*_ − (1 − *p*_*A*_), and conversely (1 − *p*_*A*_) − *p*_*A*_ for selecting the unattended patch.

We computed the value function *V* (*s*) using backwards induction over a finite horizon for each possible initial state configuration. Since trials could start with different proportions of green dots in each patch (ranging from 0.05 to 0.3 in increments of 0.01), we computed separate optimal policies for a subset of 36 combinations of initial green dot proportions in the left and right patches (6 possible proportions per patch: 0.1, 0.14, 0.18, 0.22, 0.26, 0.3). For each initial condition, we performed backwards induction over the complete state space with the following parameters: maximum number of steps = 100, maximum number of dots per patch = 100, step size (dots revealed per sampling) = 2, cost of staying = 0.005, cost of switching = 0.065, and a uniform prior (*α* = *β* = 1) for the Beta-Binomial transition function. The resulting lookup tables mapped each state to its optimal action, allowing us to simulate optimal behaviour given the reward function we defined.

The optimal value function *V*\*(*s*) was calculated recursively by iterating over all possible states and actions.

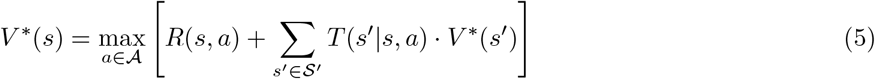

The optimal policy *π*\*(*s*) selects actions that maximize the expected sum of rewards from each state. For each state *s*, the optimal policy *π*\*(*s*) was computed using the Bellman equation:

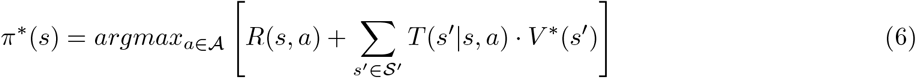

where 𝒜 is the set of possible actions, *R*(*s, a*) is the immediate reward, *T* (*s*′|*s, a*) is the transition probability to state ′, and *V* (*s*′) is the value function at the next state. 𝒮′ represents the set of possible next states given the current state and action. This yielded a lookup table mapping states to optimal actions that we used to simulate ideal observer behaviour in the task.

#### Hybrid Model

To model the choice data, we used a hybrid model that combined theory-driven and data-driven elements. The model computes action values for four possible actions: sampling from the left patch, sampling from the right patch, selecting the left patch, or selecting the right patch. The model allows for a transformation of the objective evidence into subjective magnitudes, accounting for potential effects of attention and memory. The model receives three types of objective information as input. First, the number of revealed dots (red + black) in both the attended and unattended patches. Second, the number of red dots in both patches. Third, the number of green dots in the blocked patch. The model transforms these objective quantities into subjective magnitudes through two distinct mechanisms, depending on whether it’s the first visit to a patch or a subsequent visit. During the first visit to a patch, the model accounts for potential interference from the blocked patch when estimating the unattended patch’s contents. The subjective number of dots in the unattended patch 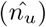 is computed as:

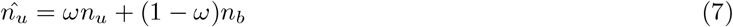

where *n*_*u*_ is the actual number of green dots in the unattended patch, *n*_*b*_ is the number of green dots in the blocked patch, and *ω* is a parameter that quantifies the resistance to interference (when *ω* = 1, there is complete resistance to interference from the blocked patch).

The model then estimates the number of red dots in the unattended patch 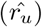 based on information from the attended patch (i.e. 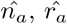):

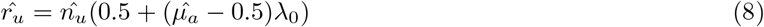

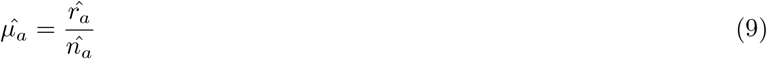

where *λ*_0_ is a free parameter that controls how strongly the proportion of red dots in the attended patch 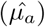 influences expectations about the unattended patch.

During the subsequent visits to a patch, information about the unattended patch must be maintained in memory rather than being directly sampled through visual perception. To capture this memory-dependent representation, the model implements an exponential decay of information about the unattended patch over time:

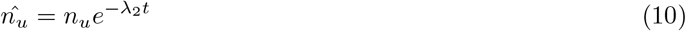

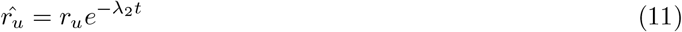

where *t* is the time elapsed since the start of the current visit and *λ*_2_ determines the rate of information decay in memory. This decay reflects the deterioration of remembered information when attention is directed elsewhere, in contrast to the perfect perception of information from the currently attended patch. Because both the remembered number of red dots 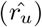 and total dots 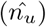 decay at the same rate, this effectively increases uncertainty about the true proportion of red dots in the unattended patch over time. In terms of Bayesian inference, this means the posterior Beta distribution over the proportion of red dots becomes increasingly wide as time passes, reflecting growing uncertainty about the unattended option.

For the attended patch, the subjective magnitudes equal the objective ones, since participants are directly looking at the patch.

Once these subjective magnitudes are computed, the model uses them to evaluate the value of each possible action. At each time point, the model considers four possible actions: selecting either patch or sampling more information from either patch. The value of selecting a patch is determined by comparing the estimated proportion of red dots between the two patches:

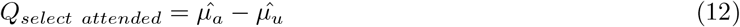

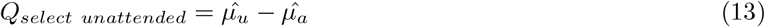

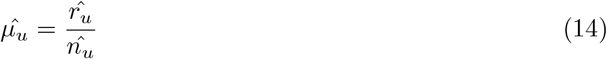

where 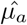 and 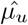 represent the estimated proportion of red dots in the attended and unattended patches, respectively.

The value of continuing to sample information is different depending on whether the participant stays with the currently attended patch or switches to the unattended patch:

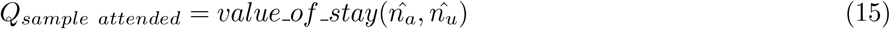

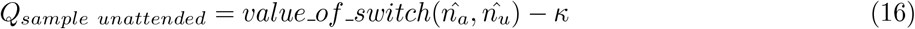

where *value of stay* and *value of switch* are functions that estimate the potential information gain from sampling, and *κ* is a free parameter that captures the cognitive cost of switching attention between patches.

To compute the value of gathering additional information (*value of stay* and *value of switch* functions), we implemented and compared three different approaches: a simple linear function, an upper confidence bound (UCB) algorithm, and an artificial neural network (ANN).

The linear function provides a straightforward relationship between the number of revealed dots and the value of sampling:

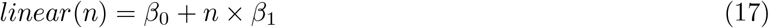

where *β*_0_ represents a baseline sampling value and *β*_1_ determines how this value scales with the number of revealed dots.

The UCB function implements a non-linear relationship between the number of revealed dots and the value of sampling:

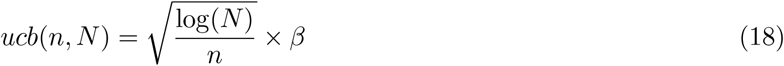

where *N* is the total number of dots across patches, *n* is the number of dots in the current patch, and *β* scales the exploration bonus. This formulation encourages sampling from patches that have been less explored relative to the total available information.

The neural network approach uses a fully connected Lipschitz-Bounded Deep Network (LBDN) with built-in guarantees on the Lipschitz bound. We implemented a network with 4 hidden layers of 32 neurons each and a tanh activation function. The network takes as input the normalized counts of revealed dots in each patch (i.e. 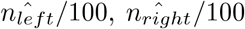), patch gaze indicators (g) [left, right], and a first-visit flag. It outputs a value of information for each patch, representing the expected utility of gathering more evidence from that location. The LBDN architecture ensures that small changes in input lead to proportionally small changes in output, providing smooth and stable value estimates.

To optimize the model parameters, we employed a multi-stage training procedure that alternates between training the neural network and the cognitive module parameters. The model was trained to minimize the cross-entropy loss between predicted and actual choices using backpropagation.

The training procedure of the hybrid model (estimate value of information using an ANN) consists of three main stages. First, we jointly optimized both the neural network and cognitive module parameters using Adam optimizer, with different learning rates for each component (*η*_*nn*_ = 0.01 for neural network, *η*_*cm*_ = 0.02 for cognitive module). This allowed both components to adapt to each other while learning at appropriate rates for their respective architectures. Next, we performed a focused optimization of the cognitive parameters using L-BFGS while keeping the neural network fixed, which helped refine the psychological parameters of the model. Finally, we conducted a fine-tuning phase where both components were again jointly optimized but with smaller learning rates (*η*_*nn*_ = 0.001, *η*_*cm*_ = 0.0001), ensuring stable convergence of the complete model. To train the standard cognitive model (linear and UCB used to compute the value of information), we used L-BFGS to minimize the cross-entropy loss between predicted and actual choices.

#### Function Recovery

To validate our hybrid modeling approach’s ability to recover underlying value-of-information computations, we conducted a series of simulation studies. We generated synthetic data from four different artificial agents, each using a distinct non-linear function to compute their value of information:

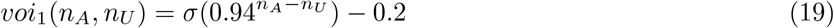

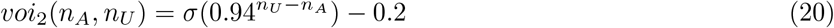

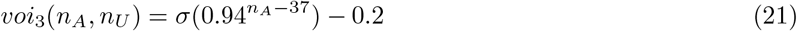

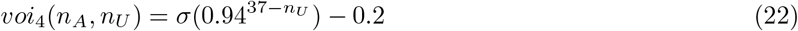

where *σ* represents the logistic function, and *n*_*A*_ and *n*_*U*_ represent the number of revealed dots in the attended and unattended patch respectively. These functions were chosen to represent different patterns of information valuation: the first two functions compute relative differences between patches, while the last two compare each patch against a fixed reference point. For each function, we simulated behavioral data using different sample sizes (4,000, 8,000, and 12,000 observations) to assess the robustness of our recovery procedure. We then applied our hybrid modeling approach to these simulated datasets, training the model to recover the underlying value-of-information function from the behavioral data alone. The recovered functions were compared to the true generating functions using Pearson correlation coefficients.

#### Feature Selection

To identify which task variables were essential for computing the value of information, we performed a systematic feature selection analysis. We first defined a full feature vector containing eight input variables: gaze position (left/right), number of dots in the blocked option, number of dots in the left patch, number of dots in the right patch, whether it was the first visit to a patch, current gaze location, value of selecting the left patch (proportion of red dots), and value of selecting the right patch. We then conducted a leave-one-out analysis where we trained separate models for each participant, comparing the full model against versions with each feature individually removed. This resulted in eight different model variants per participant. All models were trained using 8 fold cross-validation. We evaluated the impact of each feature by comparing the validation performance of the reduced models against the full input-feature model. Features were considered critical if their removal led to a significant decrease in model performance across participants.

#### Symbolic Regression

To discover interpretable mathematical equations that describe how participants compute the value of information, we performed symbolic regression analysis using the SymbolicRegression.jl package ^38^. First, we prepared the input data by binning and normalizing the number of attended and unattended dots (ranging from 0 to 100) into 24 equally spaced bins. For each unique combination of attended and unattended dots in our dataset, we computed the models predicted value of staying (continuing to sample from the currently attended patch) and switching (sampling from the unattended patch) by averaging predictions across all 20 participants fitted hybrid neural networks. We then searched for mathematical expressions that could best approximate these predicted values using symbolic regression. The search was constrained to use a limited set of mathematical operations: addition (+), subtraction (−), multiplication (), division (/), natural logarithm (log), exponential (exp), and sigmoid (*σ*) functions. To prevent excessive nesting of functions, we constrained exponential, sigmoid, and logarithmic functions to be nested at most once within functions of the same type. The complexity of constants was set to 2, and the maximum expression size was limited to 20 terms. The search process optimized for both prediction accuracy and expression simplicity, running for 10,000 iterations. The best equations were selected based on a combined score that balanced their prediction error (loss) with their complexity (computed as the number of operations and constants in the expression).

The symbolic regression discovered two core functions: a value of staying function and a value of switching function, where *N*_*attended*_ and *N*_*unattended*_ represent the number of revealed dots in the attended and unattended patches, respectively:

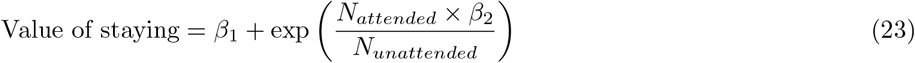

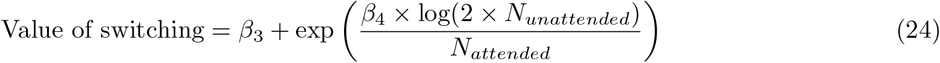

Since our feature selection analysis indicated that the first visit indicator variable was critical for optimal model performance, we allowed all parameters to vary between first visits and subsequent visits to a patch. This resulted in separate parameter sets for each phase of exploration.

However, we identified a parameter identifiability issue with the *β*_3_ parameter in the switch value function. During model fitting, both *β*_3_ (the switch function offset) and *κ*_1_ (the switching cost parameter) served as additive offsets to the overall value of switching to the unattended patch, making them mathematically non-identifiable. To resolve this issue, we reparameterized the model by removing *β*_3_ and incorporating its effect directly into the switching cost structure. The final implemented functions are:

*First visits:*

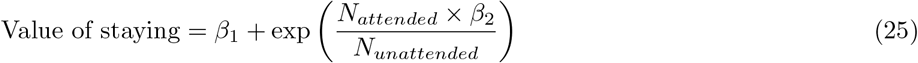

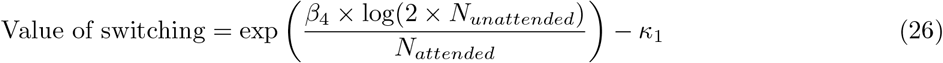

*Subsequent visits:*

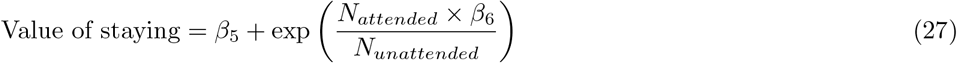

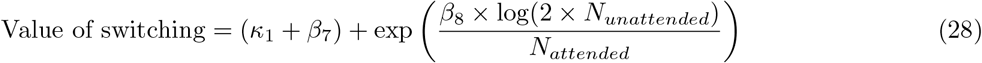

This reparameterization maintains the functional form discovered by symbolic regression while ensuring all parameters are uniquely identifiable (see Table S2).

### Fitting Symbolic and UCB Models to independent Two-Armed Bandit Task Datasets

To assess the generalizability of the insights derived from our modeling approaches, and to specifically compare the UCB model with a the Symbolic model (derived from the symbolic regression analysis), we evaluated their performance on two additional, independent datasets from a previously published study by Gershman et al. (2018; referred to as däta1änd däta2ïn our analyses, N=44 and N=44 respectively after excluding one subject from data1 that led to an extremely large value of loss when trying to fit the UCB model to their choices). These datasets involved similar information sampling tasks, providing a strong test for cross-task generalization.

For each participant in these external datasets, we fitted three models: the standard UCB model, an augmented UCB (AUCB) model, and the Symbolic model. The standard UCB model used the formulation 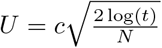, where *c* is a scaling parameter, *t* is the time step, and *N* is the number of times the option has been sampled. To control for the possibility that the Symbolic model’s superior performance was due to its additional offset parameter rather than its functional form, we also tested an augmented UCB model with the formulation 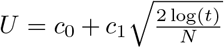, where *c*_0_ provides a baseline offset and *c*_1_ scales the exploration bonus. This AUCB model has the same parametric flexibility as the core components of our Symbolic model, allowing for a more controlled comparison of functional forms.

To evaluate how well each model could predict each participant’s choices, we employed a Leave-One-Out (LOO) cross-validation procedure on a per-subject basis. This involved iteratively training each model on all but one trial for a given subject and then testing its predictive accuracy on the held-out trial. This process was repeated such that every trial served as part of a test set.

The primary metric for evaluating model performance was the average prediction loss (e.g., cross-entropy loss) on these held-out test trials, calculated for each subject for both models. This provides a measure of how well each model generalized to unseen data within each participant from the external datasets.

The per-subject average LOO losses for the Symbolic model and the UCB model were then statistically compared to determine if one model offered consistently better predictions of choice behavior on these novel datasets. This comparison involved paired t-tests on the differences in mean losses per subject, as visualized in the comparative loss plot.

### Imaging data acquisition

Structural and functional MRI data was collected with a Siemens 7 Tesla MRI scanner. High-resolution functional data were acquired with a multiband gradient echo T2* echo planar imaging sequence with 1mm isotropic voxels, multiband acceleration factor 2, repetition time (TR) = 1.378s, echo time (TE) = 27ms, flip angle = 90°, and GRAPPA acceleration factor 2. The parameters were selected to maximise signal-to-noise ratio in subcortical areas. To accommodate the high temporal and spatial resolution of the protocol, functional scans had a limited field of view (FOV) oriented at 45 degrees with respect to the AC-PC line (36 slices). The FOV captured all regions of interest in the midbrain, brainstem and cortex. Before acquiring the task-related functional scan, we acquired a presaturation single-measurement, whole-brain functional scan with the same orientation. The pre-saturation scan was used to facilitate registration of the limitedFOV task-related functional scan to the whole brain. Structural data were acquired using a T1-wieghted MP-RAGE sequence with 0.7mm isotropic voxels, GRAPPA acceleration factor 2, TR = 2200ms, TE = 3.02ms, and; inversion time (TI) = 1050ms. To correct distortions arising from inhomogeneities in the magnetic field, a fieldmap sequence was acquired with 2mm isotropic voxels, TR = 620ms, TE1 = 4.08ms, and TE2 = 5.1ms. To account for the effects of physiological noise on functional MRI data, participants were fitted with a pulse oximeter and respiratory bellows that acquired cardiac and respiratory timeseries at 50Hz using a BioPac MP160 device (BIOPAC Systems Inc., USA).

### fMRI data preprocessing

Preprocessing of fMRI data was performed with the FMRIB Software Library ^65,66^. The Brain Extraction Tool ^67^ was used to separate brain from non-brain matter in structural and functional images. Functional images were normalised, spatially smoothed (Gaussian kernel with a 3mm full-width half-maximum) and temporally high-pass filtered (3 dB cut-off = 100s), and artefacts arising from head motion were removed using MCFLIRT^68^. Registration of task-related functional images to Montreal Neurological Institute (MNI)-space was performed in three stages: 1. The task-related limited-FOV EPI was registered to the pre-saturation whole-brain EPI using FMRIB’s Linear Image Registration Tool with 6 degrees of freedom transformation. 2. The whole-brain EPI was registered to the subject-specific structural images using Boundary-Based Registration (BBR) incorporating fieldmap correction ^69^. 3. Subject-specific structural images were registered to a 1mm resolution Standard MNI template with FMRIB’s Non-linear Registration Tool (FNIRT; ^65^).

### fMRI data analysis

Statistical analysis of whole-brain functional data was performed at three levels using FMRIB’s Expert Analysis Tool (FEAT; ^65,66^). In the first level, a univariate general linear model was used to compute parameter estimates for each regressor in each session ^70^. Contrast and variance estimate for each parameter in each participant were subsequently combined in a fixed-effects analysis conducted at the second level. Finally, a random-effects analysis was conducted at the third level, where subject-identity was a random effect ^71^. Significance testing was performed with cluster-correction, a cluster significance threshold of *P* = .001, and a voxel inclusion threshold of *z* = 3.1. Data were pre-whitened before analysis to account for temporal autocorrelations in BOLD signal. We performed one-whole brain analyses for each of the three different approaches to computing the value of information. The GLM identified voxels where BOLD signal represented the value of staying, switching, or selecting the attended or unattended patch.

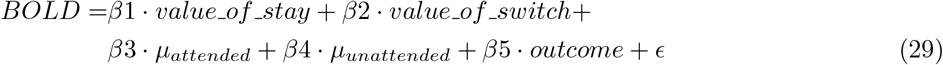

All regressors were convolved with a double-gamma hemodynamic response function. Further non-task confound regressors were added to reduce noise in BOLD signal, including: (1) head motion parameters estimated using MCFLIRT during pre-processing ^68^; (2) regressors for voxel-wise estimates of physiological noise arising from cardiac and respiratory activity, estimated using FSL’s Physiological Noise Monitoring (PNM) tool ^72^, and; (3) regressors for motion outliers, indicating volumes with head motion that could not be corrected with linear methods.

To compare the three different approaches to computing the value of information (linear, UCB, and ANN), we ran identical GLM analyses for each approach, varying only in how the value of staying and switching were computed. For each of the 80 sessions (4 sessions X 20 subjects), we fitted three separate GLMs using the value computations from each model. We then computed the Mean Squared Error (MSE) between the GLM predictions and the actual BOLD signal for each session, resulting in 80 MSE values per model. To statistically compare the performance of the three approaches, we conducted non-parametric Wilcoxon signed-rank tests on these paired MSE values, allowing us to assess whether one approach consistently provided better predictions of neural activity than the others.

### ROI analysis

To analyze activity in specific regions of interest (ROIs: VTA, SN, DRN, LC, VSN), we extracted voxel-wise beta estimates (effect sizes) and their corresponding standard errors from the first-level GLM analysis for each relevant contrast (contrast of parameter estimates, or “cope” in FSL). For each ROI and contrast, we fitted a linear mixed-effects model using the MixedModels.jl package in Julia to estimate the group-level effect. The model formula was *effsize* ~ 1 + (1|*subject*) + (1|*voxel*), predicting the voxel-wise beta estimate (effsize) with a fixed intercept (representing the group effect) and random intercepts for subject and voxel to account for inter-subject and inter-voxel variability. Crucially, these models were weighted by the inverse variance of the beta estimates (1*/standard error*^2^) to give more influence to more precise measurements at the voxel level. This approach yields a robust estimate of the average activation (the fixed intercept) for each contrast within each ROI across the group, while appropriately accounting for different sources of variance. P-values for the fixed intercept were extracted and corrected for multiple comparisons across the tested ROIs using the Bonferroni method.

### RSA analysis

Representational Similarity Analysis (RSA) was employed to test whether the similarity structure of neural activity patterns related to the similarity structure of behavioral sampling duration on a trial-by-trial basis within each session. For each participant and session, we constructed two types of Representational Dissimilarity Matrices (RDMs). First, a behavioral RDM was computed based on the number of samples taken in each trial; the dissimilarity between any pair of trials (i, j) was defined as the Euclidean distance between their respective sample counts. Second, neural RDMs were constructed for specific regions of interest (VTA, SN, AI, ACC). To control for ROI size and shape differences, we used masks matched to the VTA’s shape, positioned within each target region. Trial-level BOLD activation patterns (z-statistics from the first-level GLM) were extracted for each trial within these masks. The dissimilarity between neural patterns for any pair of trials (i, j) was calculated as the Euclidean distance between their multi-voxel activation vectors. Both behavioral and neural RDMs were then vectorized by taking their upper triangular elements (excluding the diagonal). The correspondence between neural patterns and behavior was quantified by computing the Kendall’s *τ* correlation between the vectorized neural RDM and the vectorized behavioral RDM for each session. These session-level correlation coefficients were averaged within each participant for each ROI. Group-level significance was assessed using a one-sample Wilcoxon Signed Rank Test against zero across participants for each ROI, testing for positive correlations. P-values were Bonferroni corrected for the number of ROIs tested.

