## Supplemental Materials for "Interpretable abstractions of artificial neural networks predict behavior and neural activity during human information gathering"

### Supplementary Figures

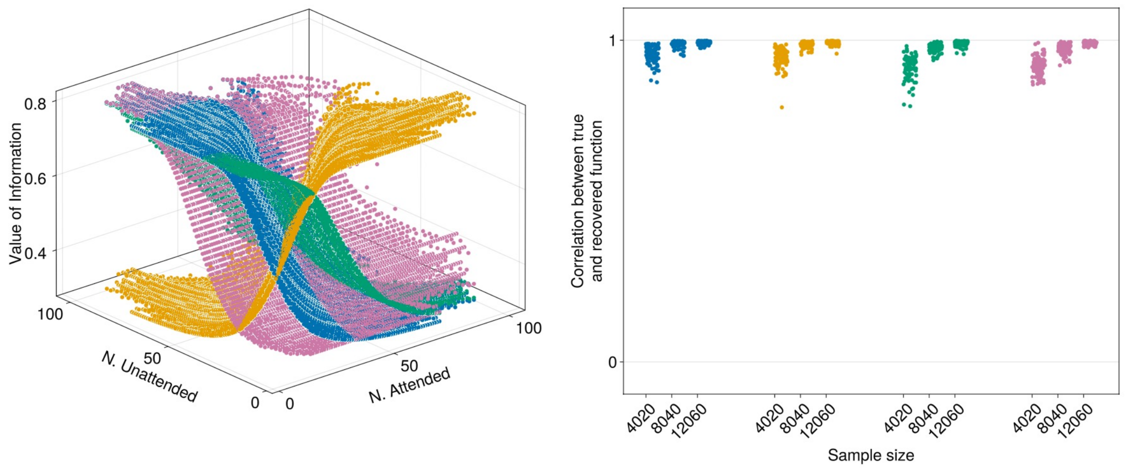


**Supplementary Fig. 1** Function recovery validation. Left: Three-dimensional scatter plot showing the value of information as a function of the number of dots in the attended (N. Attended) and unattended (N. Unattended) patches. Different colors (blue, green, pink, and orange) represent different simulated functions used to generate the data. Right: Scatter plot showing the correlation between true and recovered functions across different sample sizes (4020, 8040, and 12060 observations). Each color represents a different simulated function, with each dot representing a single recovery attempt.

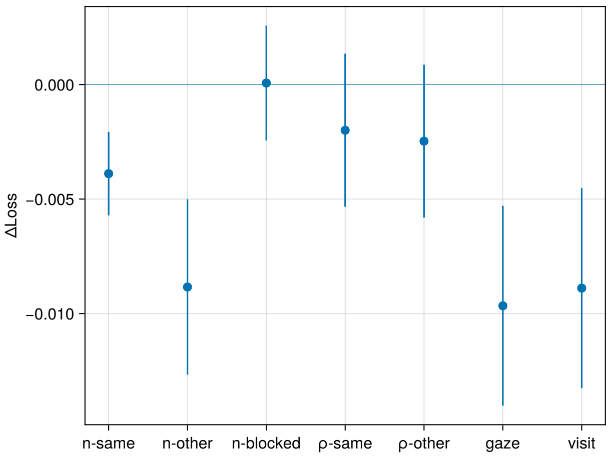


**Supplementary Fig. 2** Feature importance analysis. Dot plot showing the change in model loss (ΔLoss) when different features are removed from the model. Each blue dot represents the mean loss difference, with vertical blue lines indicating confidence intervals. Features tested include n-same (number of dots in the same patch), n-other (number of dots in the other patch), n-blocked (number of dots in the blocked patch), $\rho$-same (proportion of red dots in the same patch), $\rho$-other (proportion of red dots in the other patch), gaze (current gaze position), and visit (whether it's the first visit to a patch).

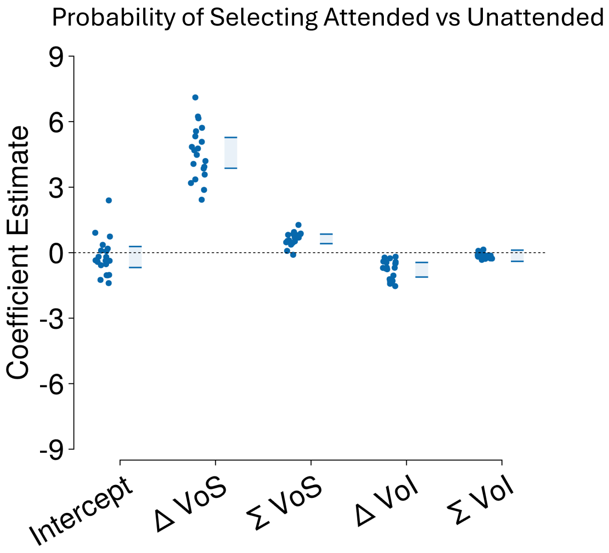


**Supplementary Fig. 3** Selection behavior analysis. Regression analysis predicting the probability of selecting the attended versus unattended patch. Left: Coefficient estimates from a mixed-effects logistic regression model. Each light blue dot represents a subject-specific random effect estimate, with light blue rectangles showing the ±95% confidence intervals of the fixed effects. Predictors include the intercept, value of selecting the attended option (VoS attended), value of selecting the unattended option (VoS unattended), value of information for the attended option (VoI attended), and value of information for the unattended option (VoI unattended). Right: AIC comparison between models using different value of information computations (Linear, UCB, and ANN), with lower values indicating better fit.

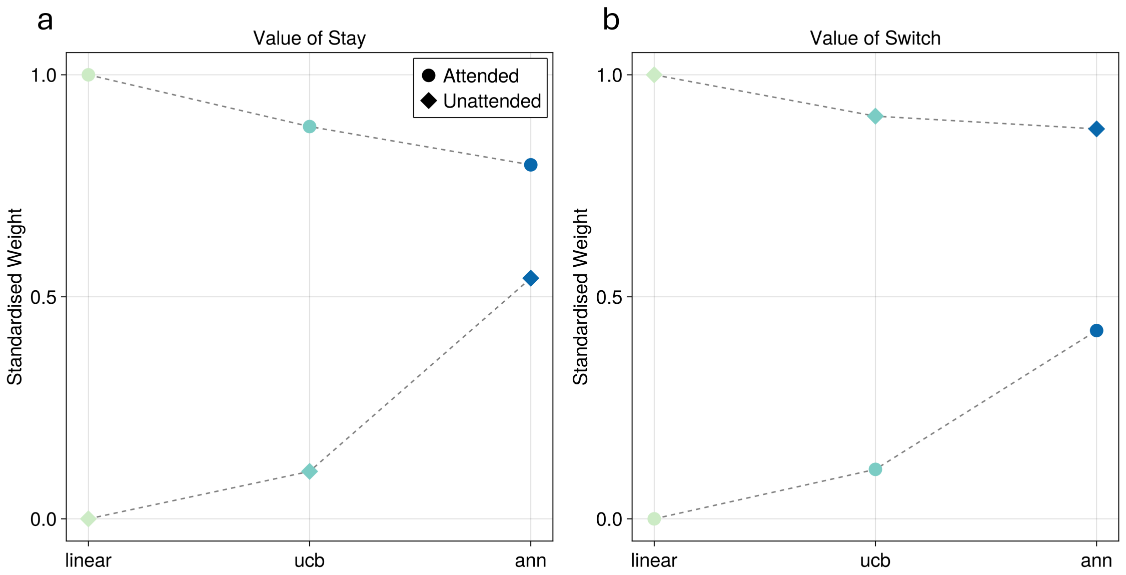


**Supplementary Fig. 4** Model comparison for evidence weighting. a, Standardized weights for the value of stay computation across three models (linear, UCB, and ANN). Circles represent weights for the attended evidence, while diamonds represent weights for the unattended evidence. Colors transition from light green to dark blue across models. Dotted lines connect the weights across models. b, Standardized weights for the value of switch computation across the same three models, using the same visual conventions as in panel a.

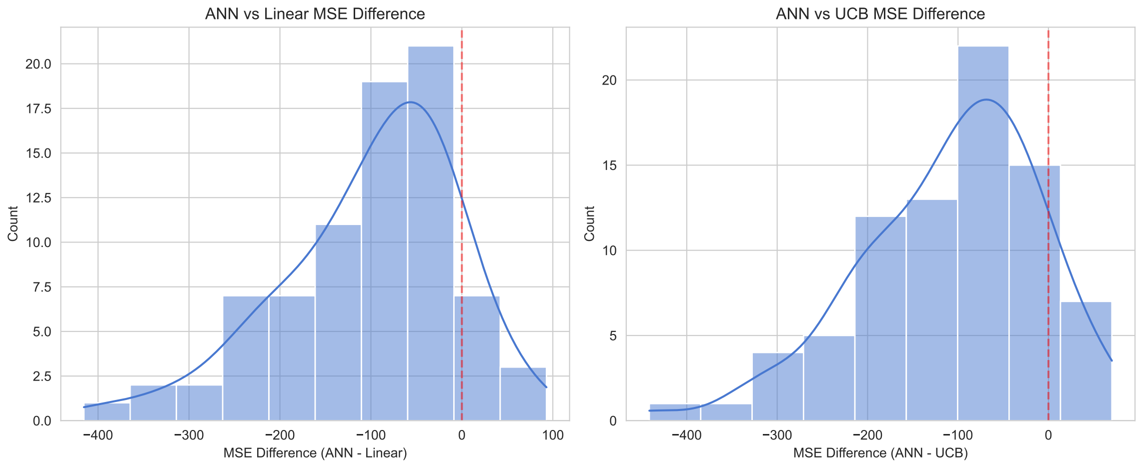


**Supplementary Fig. 5** Neural model comparison. Histograms showing the distribution of Mean Squared Error (MSE) differences between models in predicting neural activity. Left: Histogram of MSE differences between the ANN and Linear models (ANN - Linear), with negative values indicating better performance by the ANN model. Right: Histogram of MSE differences between the ANN and UCB models (ANN - UCB). Both histograms include a blue curve showing the estimated probability density and a vertical red dashed line at zero.

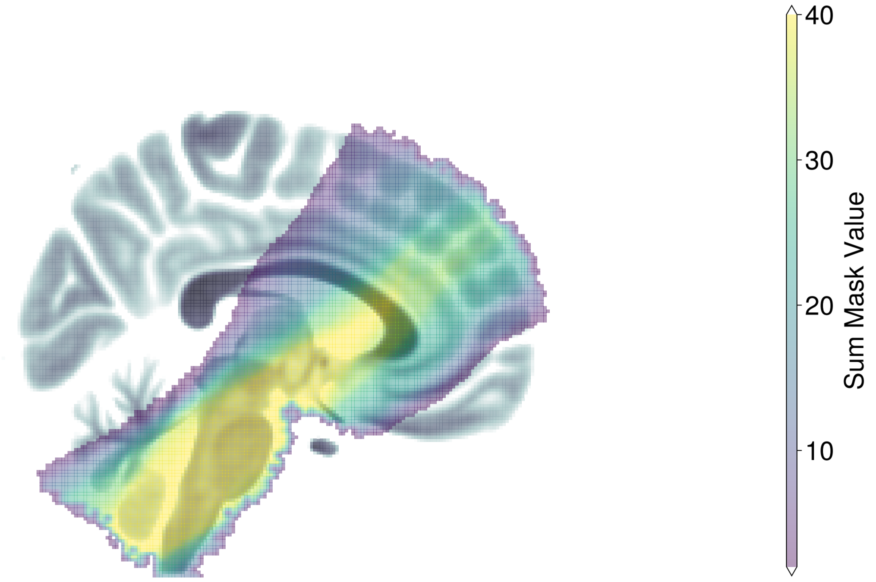


**Supplementary Fig. 6** Field of view coverage across sessions. Sagittal view (x = 87) showing the spatial coverage of the limited field of view functional imaging across all 40 sessions (2 sessions × 20 participants). The grayscale background shows the MNI152 T1 1mm brain template. The colored overlay represents the sum of individual session masks, with the color scale indicating the number of sessions with coverage at each voxel location. Warmer colors indicate regions consistently captured across more sessions, while cooler colors represent areas with coverage in fewer sessions.

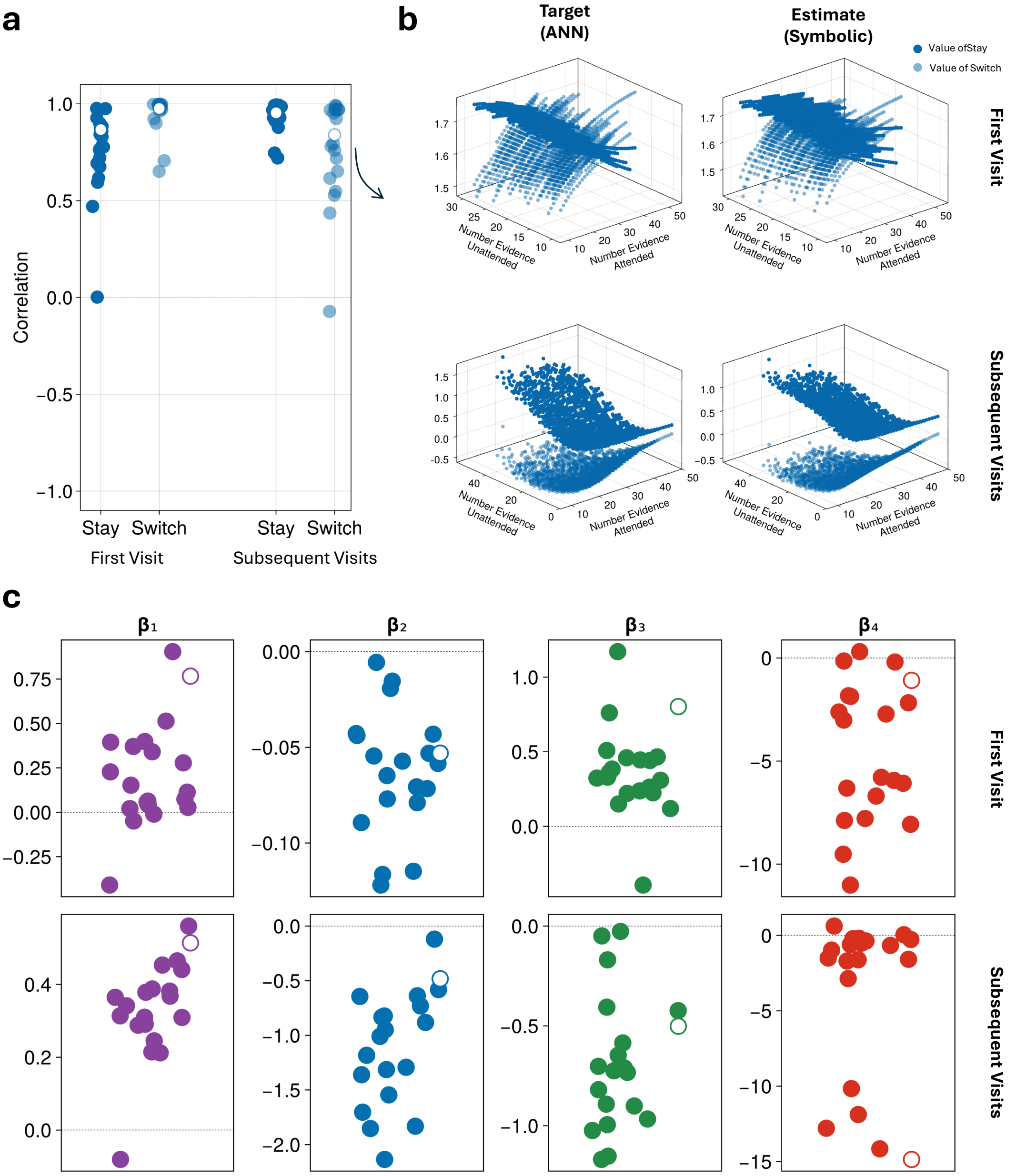


**Supplementary Fig. 7** Symbolic regression approximation of ANN-derived value of information functions. a. Correlation between ANN-generated value of information and symbolic function approximations across all participants (n=20). Four correlations are shown for value of staying and switching during first visits and subsequent visits to patches. Each point represents one participant, with the highlighted participant (white circle) shown in detail in panel b. b. Example participant showing 3D visualization of value of information functions. Left column shows target functions from the trained ANN model; right column shows estimates from the optimized symbolic functions. Top row displays first visit functions, bottom row shows subsequent visit functions. Dark points represent value of staying, light points represent value of switching. c. Individual parameter estimates ($\beta_1 - \beta_8$) controlling the symbolic functions across all participants. Each parameter corresponds to specific components of the value of information computation: $\beta_1$,$\beta_5$ (stay VOI offsets), $\beta_2$,$\beta_6$ (stay VOI scaling), $\beta_3$,$\beta_7$ (switch VOI offsets), $\beta_4$,$\beta_8$ (switch VOI scaling) for first and subsequent visits respectively. The highlighted participant (white circle) corresponds to the example shown in panel b.

### Supplementary Tables

**Table S1** Comparison of ANN vs LINEAR and ANN vs UCB models across brain regions

| ROI | ANN vs LINEAR | | ANN vs UCB | |
|---|---|---|---|---|
| | Median Diff. | p-value* | Median Diff. | p-value* |
| VTA | −31.12 | 0.009 | −37.63 | 0.003 |
| SN | −18.626 | $6.05 \times 10^{-4}$ | −18.556 | $3.33 \times 10^{-4}$ |
| LC | −67.461 | 0.041 | −87.452 | 0.083 |
| DRN | −84.7 | $4.50 \times 10^{-4}$ | −90.77 | 0.002 |
| VSN | −110.112 | $9.00 \times 10^{-6}$ | −117.639 | $1.34 \times 10^{-6}$ |

*Bonferroni corrected p-values. All comparisons based on n=80.
Negative values indicate ANN model performed better (lower MSE).

**Table S2** Model parameters and descriptions

| Model | Parameter | Count | Description |
|---|---|---|---|
| **All Models** | $\lambda_0$ | 1 | **First-visit proportion estimate**: Controls how the proportion of red dots observed in the attended patch influences the estimate of red dots in the unattended patch during first visit. $\lambda_0 = 0$ assumes 50% red dots, $\lambda_0 = 1$ fully uses attended patch proportion. |
| | $\omega$ | 1 | **Interference resistance**: Resistance to interference from blocked/irrelevant options. Values closer to 1 indicate better ability to ignore task-irrelevant information. |
| | $\kappa_1$ | 1 | **Switching cost**: Cost of switching attention between patches. Higher values make attention switches more costly, promoting sustained attention. |
| | $\lambda_2$ | 1 | **Memory decay rate**: Rate at which information about unattended patches decays in memory over time. Higher values lead to faster forgetting. |
| | $\tau$ | 1 | **Decision temperature**: Controls the stochasticity of action selection. Lower values make choices more deterministic, higher values more random. |
| **Linear** | $\beta_1$ | 1 | **VOI offset**: Baseline value of information, independent of evidence collected. Sets the overall propensity to sample information. |
| | $\beta_2$ | 1 | **VOI slope**: Linear scaling factor determining how value of information changes with amount of evidence. Negative values create diminishing returns. |
| | **Total** | **7** | **Linear relationship**: VOI $= \beta_1 + \beta_2 \times evidence$ |
| **UCB** | $\beta$ | 1 | **Exploration bonus**: Scaling factor for the Upper Confidence Bound exploration bonus. Controls the strength of the uncertainty-driven exploration. |
| | **Total** | **6** | **Non-linear relationship**: VOI $= \beta \times \sqrt{\log(N)/n}$, where $N$ is total evidence and $n$ is evidence from current patch |
| **Symbolic** | $\beta_1$ | 1 | **Stay VOI offset (first visit)**: Baseline value for continuing to sample from current patch on first visit. |
| | $\beta_2$ | 1 | **Stay VOI scaling (first visit)**: Scaling parameter for evidence interaction in stay computation during first visit. |
| | $\beta_3$ | 1 | **Switch VOI offset (first visit)**: Baseline value for switching to alternative patch on first visit. |
| | $\beta_4$ | 1 | **Switch VOI scaling (first visit)**: Scaling parameter for evidence ratio in switch computation during first visit. |
| | $\beta_5$ | 1 | **Stay VOI offset (after first visit)**: Baseline value for continuing to sample from current patch after first visit. |
| | $\beta_6$ | 1 | **Stay VOI scaling (after first visit)**: Scaling parameter for evidence interaction in stay computation after first visit. |
| | $\beta_7$ | 1 | **Switch VOI offset (after first visit)**: Baseline value for switching to alternative patch after first visit. |
| | $\beta_8$ | 1 | **Switch VOI scaling (after first visit)**: Scaling parameter for evidence ratio in switch computation after first visit. |
| | **Total** | **13** | **Discovered functions**: Stay $= \beta_1 + \exp((N_{attended} \times \beta_2)/N_{unattended})$, Switch $= \beta_3 + \exp(\beta_4 \times \log(2 \times N_{attended})/N_{unattended})$ |
| **ANN** | $\theta$ | 7,592 | **Neural network weights**: Parameters of the Lipschitz-Bounded Deep Network (4 hidden layers × 32 neurons each). Learns complex non-linear mappings from task state to value of information. |
| | **Total** | **7,597** | **Data-driven relationship**: VOI = LBDN($N_{attended}/100$, $N_{unattended}/100$, gaze_position, first_visit) |
